# The annexin A1 mimetic peptide Ac2-26 promotes inflammation-resolution and stimulates skeletal muscle regeneration in the *mdx* mouse model of Duchenne muscular dystrophy

**DOI:** 10.64898/2026.10.05.756885

**Authors:** Hamood Rehman, Xinyue Lu, Savannah M Nasrallah, Binayok Sharma, Sophie Lies, Shikha Adhikari, Steven S. Welc, James F. Markworth

**Affiliations:** Department of Animal Sciences, College of Agriculture, Purdue University, West Lafayette, IN, USA; Interdepartmental Nutrition Program (INP), College of Health and Human Sciences, Purdue University, West Lafayette, IN, USA; Center for Aging and the Life Course (CALC), Purdue University, West Lafayette, IN, USA; Department of Anatomy, Cell Biology & Physiology, Indiana University School of Medicine, Indianapolis, IN; Indiana Center for Musculoskeletal Health (ICMH), Indiana University, Indianapolis, IN, USA; Purdue Institute of Inflammation, Immunology, and Infectious Disease (PI4D), Purdue University, West Lafayette, IN, USA

## Abstract

Duchenne muscular dystrophy (DMD) is an X-linked disorder mainly affecting young boys characterized by repeated cycles of myofiber injury/regeneration and chronic muscle inflammation. Although glucocorticoids can slow disease progression, their potential long-term adverse effects highlight the need for new steroid-sparing therapies for DMD. Annexin A1 (ANXA1) is a glucocorticoid-induced pro-resolving protein that signals primarily through formyl peptide receptor 2 (FPR2). We investigated whether the ANXA1 mimetic peptide Ac2-26 could reduce inflammation, stimulate tissue regeneration, and improve skeletal muscle function in muscular dystrophy. 3-week-old dystrophin-deficient male *mdx* mice were randomized to receive daily intraperitoneal injections of vehicle control, deflazacort (1 mg/kg/day), or Ac2-26 (1 mg/kg/day) for a duration of 5-weeks. Body composition, skeletal muscle mass, contractile function, and tissue histopathology/immune-cell infiltration were assessed at 8-weeks of age. Direct effects on myogenesis were also examined *in vitro* using primary D2-*mdx* myoblasts. ANXA1 was expressed predominantly by myeloid cells rather than resident dystrophic muscle cells. Both deflazacort and Ac2-26 markedly reduced intramuscular inflammation and increased myofiber size. However, Ac2-26 produced broader anabolic and functional benefits, increasing body-weight gain, lean body mass, hindlimb muscle weights, and absolute contractile force. Ac2-26 treatment also directly promoted myogenesis of dystrophic myoblasts *in vitro*, while pharmacological FPR2 blockade impaired myotube development. Overall, Ac2-26 mimicked the anti-inflammatory effects of deflazacort while providing superior benefits for body weight and composition, skeletal muscle growth, myofiber regeneration, and absolute muscle strength. These findings support therapeutic targeting of the ANXA1–FPR2 axis as a potential safe and effective steroid-sparing immunomodulatory strategy for DMD.

## INTRODUCTION

Duchenne muscular dystrophy (DMD) is a severe X-linked neuromuscular disorder^1^. DMD affects 1 in 3,500^2^ to 1 in 5,000^3^ live male births globally. It is caused by mutations in the *DMD* gene that either greatly reduces or totally eliminates production of the dystrophin protein^4^. Dystrophin helps stabilize the sarcolemma during muscle contractions, so its absence leaves muscle cells vulnerable to repeated membrane damage, necrosis, and incomplete healing^5,6^. Over time, this process leads to worsening muscle weakness, loss of ambulatory ability, cardiomyopathy, respiratory failure, and early death^7–9^. While restoring functional dystrophin expression remains the main treatment focus, inflammation and fibrosis pathways also play significant roles in disease progression and are important targets for additional therapies.

Much of the mechanistic research in DMD relies on the *mdx* mouse which harbors a spontaneously genetic mutation resulting in the absence of dystrophin^1^. While the dystrophic phenotype of the *mdx* mouse is generally less severe than human DMD in several limb muscles, it still exhibits key disease features like repeated muscle degeneration and regeneration, immune-cell infiltration, and ongoing inflammatory signals^10–14^.

In dystrophic muscle, inflammation isn’t just a byproduct of fiber damage; it plays a crucial role in amplifying injury^15,16^. Polymorphonuclear neutrophils (PMNs) arrive early following muscle injury and can directly worsen myofiber damage^17,18^. Depletion of PMNs from host *mdx* mice has been found to reduce the necrosis in dystrophic muscle^19^. Blood monocytes and monocyte-derived tissue macrophages (MΦ) add further complexity. Antibody mediated MΦ depletion has been shown to result in a >75% reduction in injured fiber concentration compared with nondepleted *mdx* mice^20^. Specifically, M1-like pro-inflammatory MΦ dominate the inflammatory infiltrate of dystrophic muscle during peak necrosis and promote tissue harm through cytotoxic mediators and cytokines^21^. In contrast, M2-like MΦ help clear debris, support muscle formation, and assist repair^22^. If the transition from M1 to M2 MΦ is delayed or incomplete, inflammation persists and tends to lead to fibro-adipogenic changes and fibrosis, hindering proper regeneration^23,24^.

Glucocorticoids have been used as anti-inflammatory and immunosuppressive drugs for years^25^. They are believed to block inflammation, in part, by inhibiting the phospholipase A_2_ (PLA_2_) which catalyzes the release of the omega-6 (n-6) long chain polyunsaturated fatty acid (LC-PUFA) arachidonic acid (20:4n-6) from cell membrane phospholipids^26^. A reduced intracellular availability of free intracellular ARA limits the downstream production of pro-inflammatory lipid mediators including prostaglandins and leukotrienes^27^. Synthetic glucocorticoid drugs including prednisone, prednisolone, and deflazacort have each been shown to slow the decline in function and help maintain muscle strength in human DMD patients^28,29^. Deflazacort is often chosen as an alternative to prednisone because it offers similar benefits and may cause less weight gain in some cases^30,31^. However, long-term use of glucocorticoids involves significant side effects such as growth suppression, weight gain, weakened bones, hormonal and metabolic issues, behavioral changes, and other toxic effects, which limit their prolonged use in children^32–34^. Importantly, the exact molecular pathways through which steroids benefit dystrophic skeletal muscles are not yet fully understood. An improved understanding of these pathways could lead to steroid-sparing treatments that provide the same benefits without the extensive side effects of long-term steroid therapy.

Annexin A1 (ANXA1) is a phospholipid-binding protein regulated by glucocorticoids that has anti-inflammatory and pro-resolving roles in the innate immune response^35–39^. Glucocorticoids can enhance ANXA1 expression in myeloid cells and other systems, driving its gene transcription and/or extracellular release of pre-formed stores^40,41^. Once outside the cell, ANXA1 and its N-terminal peptides primarily signal via formyl peptide receptor 2 (FPR2/ALX), a G protein–coupled receptor found on the cell surface of blood leukocytes and other cell types^42,43^. ANXA1–FPR2 signaling attenuates early stages of the acute inflammatory response by restricting PMN–endothelial transmigration and promoting PMN apoptosis^44^. Furthermore, ANXA1 plays an active role in the resolution phase of the acute inflammatory response by enhancing blood monocyte recruitment to the site of inflammation^45^ and facilitating MΦ-mediated clearance of apoptotic neutrophils (efferocytosis)^46^. In addition to its signaling through extracellular receptors, annexin proteins including ANXA1 are involved in intracellular calcium-dependent membrane remodeling and repair^47^. Beyond its immunomodulatory actions ANXA1 is also essential for myoblast migration^48^, muscle cell fusion^49^, and skeletal muscle regeneration following acute injury^46^. Interestingly, myeloid cell specific knockout of ANXA1 has similar deleterious effect on muscle regenerative capacity following acute muscle injury as whole-body ANXA deficiency, suggesting that infiltrating blood leukocytes rather than resident muscle cells are a major cellular source of ANXA1 in injured tissue^50^. These combined pro-resolving and pro-regenerative functions make ANXA1 a promising candidate for potentially mediating, in part, the beneficial effects of glucocorticoid therapy in dystrophic muscle.

Consistent with the role of ANXA1 in limiting inflammation and promoting tissue regeneration, systemic administration of recombinant ANXA1 has been found to reduce tissue PMN influx and/or promote MΦ-mediated efferocytosis of PMNs in many acute injury models^51–54^. The N-terminal ANXA1 mimetic peptide Ac2-26 offers a practical approach for testing this pathway therapeutically. It replicates many of the anti-inflammatory and pro-resolving functions of the parent protein and has demonstrated protective effects in various models of sterile injury and inflammation^51,55,56^. Recent studies indicate that systemic injection of mice with Ac2-26 limits neutrophil infiltration in skeletal muscle after acute ischemia-reperfusion injury^57^ and provides cardioprotective benefits following a myocardial infarction^58^. Another study suggested that Ac2-26 induces skeletal muscle cell migration in *in vitro* C2C12 cell culture model^48^. Consequently, Ac2-26 could serve as a novel steroid-sparing treatment to decrease chronic muscle inflammation and encourage myofiber repair in patients with DMD. Pharmacologic inhibitors are also useful for exploring pathway dependence: Boc-2, a broad-spectrum formyl peptide receptor inhibitor, and WRW4, which more specifically targets FPR2, can be employed to determine if effects driven by ANXA1 or Ac2-26 in muscle rely on FPR2^59,60^.

In the current study, we tested the effect of long-term daily treatment with the ANXA1 mimetic peptide Ac2-26 (1 mg/kg/day) when compared to deflazacort (1 mg/kg/day) as the current standard of care in the *mdx* mouse model of DMD. Both deflazacort and Ac2-26 greatly reduced skeletal muscle inflammation and increased myofiber size in the fast-twitch TA muscle. However, only Ac2-26 increased absolute muscle mass and strength. In contrast, deflazacort negatively impacted weight gain and lean body mass. Treatment with Ac2-26, but not deflazacort, also stimulated *in vitro* myogenesis in D2-*mdx* myoblasts via an FPR2. These results show that targeting the ANXA1/FPR2 pathway with ANXA1 mimetic peptides can promote inflammation-resolution and skeletal muscle regeneration in *mdx* mice, highlighting this axis as a potential steroid-sparing therapy for DMD.

## MATERIALS AND METHODS

### Animals and treatments

Male C57BL/10ScSn-*Dmd^mdx^*/J (*mdx*) mice were obtained from The Jackson Laboratory (Strain #001801), and a breeding colony was established in-house. Mice were maintained with *ad libitum* access to food and water under controlled housing conditions. At 23 days of age *mdx* mice were randomly assigned to various treatment groups. In daily administration protocols, animals received once-daily intraperitoneal (IP) injections of sterile saline (13.33 µL/gram body weight) containing vehicle control (0.25% DMSO; n = 10), deflazacort (1 mg/kg/day; n = 10)^61^, or Ac2-26 (1 mg/kg/day; n = 11)^62–65^ for a duration of 5 weeks^61^. In weekly administration protocols, animals received once-weekly IP injections of sterile saline (13.33 µL/gram body weight) containing vehicle control (0.25% DMSO; n = 8), deflazacort (1 mg/kg/week; n=10), or Ac2-26 (1 mg/kg/week; n = 8) for a duration of 5 weeks. Body weights were recorded daily throughout the treatment period to monitor longitudinal changes. At 8-weeks of age body composition was assessed using magnetic resonance-based analysis (EchoMRI 3-in-1) to quantify lean and fat mass.

### Muscle force testing

Contractile function of hind limb muscles was evaluated both *in vivo* and *in situ*. For *in vivo* measurements, mice were anesthetized with 2% isoflurane one day prior to euthanasia and positioned on a heated platform maintained at 37 °C. The ankle joint plantar flexor muscle group (gastrocnemius, plantaris, and soleus) of the right hindlimb was tested using a non-invasive system (Aurora Scientific, 1300A 3-in-1). The limb was stabilized by securing the knee, while the foot was affixed to a footplate. Needle electrodes were placed over the posterior lower hindlimb skin overlying the gastrocnemius muscle, and stimulation current was optimized to obtain maximal isometric contractions. Peak isometric twitch and tetanus torque measurements were recorded, followed by assessment of the force–frequency relationship using stepwise increases in stimulation frequency including 10, 25, 50, 80, 100, 150, 200, 300 Hz with one minute rest intervals between contractions. Animals were allowed to recover overnight prior to *in situ* muscle force testing^66^.

Terminal *in situ* measurements were performed on the day of euthanasia. Mice were anesthetized with 2% isoflurane, the right tibialis anterior (TA) muscle was carefully exposed, and its distal tendon was freed and attached to a force transducer lever using a 4-0 silk suture (Aurora Scientific, 1300A 3-in-1). The knee was pinned to maintain a fixed position during testing and mice were positioned on a heated platform maintained at 37 °C. Muscle activation was achieved via electrodes positioned between the TA muscle and the overlying fascia. Stimulation current and muscle length were adjusted to determine the optimal length (*L*_0_), at which maximal twitch force (*P*_t_) was obtained. Tetanic contractions were then induced at a stimulation frequency of 150 Hz to determine maximal tetanic force (*P*_0_), and the force-frequency relationship established using stepwise increases in stimulation frequency including 10, 25, 50, 80, 100, 150, 200, 300 Hz with one minute rest between contractions to minimize fatigue. The exposed muscle was continuously moistened with phosphate-buffered saline (PBS) to prevent dehydration^66^.

### Tissue collection

After *in situ* muscle contractile testing, mice were euthanized under isoflurane anesthesia via cervical dislocation and bilateral pneumothorax. Hindlimb muscles, including the gastrocnemius (GAST), plantaris (PLA), soleus (SOL), tibialis anterior (TA), and extensor digitorum longus (EDL), were carefully dissected from both hindlimbs and weighed. The heart was also collected and weighed. The distal half of the TA, EDL, PLA, SOL, and GAST, as well as the left ventricle of the heart was oriented on plastic support, embedded in Optimal Cutting Temperature (OCT) compound, and frozen in isopentane precooled in liquid nitrogen for histological analysis. The remaining tissues were snap-frozen in liquid nitrogen. All samples were stored at −80°C until further analysis.

### Immunohistochemistry

Immunohistochemistry was performed using the method previously described^67^. Tissue cross-sections (10 µm) were cut from the mid-belly region of OCT-embedded TA, SOL, and left ventricle of the heart using a Leica CM1950 cryostat at −20°C. Sections were collected onto SuperFrost Plus slides and air-dried at room temperature. Unfixed sections were used for hematoxylin and eosin (H&E) and muscle fiber type staining. Tissue sections designated for immune cell staining were fixed in acetone for 10 minutes at −20°C and then air-dried prior to staining.

Slides were blocked using either 10% normal goat serum (GS) (Invitrogen, 10000C) in PBS or Mouse-on-Mouse (M.O.M.) IgG Blocking Reagent (Vector Laboratories, MKB22131) whenever mouse primary antibodies were used. Sections were then incubated overnight at 4°C with primary antibodies diluted in either 10% GS in PBS or M.O.M. protein diluent. The following day, slides were washed in PBS and incubated with specific Alexa Fluor– conjugated secondary antibodies (1:500 in PBS) for 1 h at room temperature. After washing, sections were mounted using MOWIOL fluorescence mounting medium, and nuclei were counterstained with DAPI (Invitrogen, D21490, 2 μg/mL).

Primary antibodies used included ANXA1 (Abcam, ab214486, 1:500), Ly6G (Bio-Rad, MCA6077GA, 1:50), CD68 (Bio-Rad, MCA1957, 1:200), CD206 (Bio-Rad, MCA2387, 1:50), MyHC type I (DSHB, BA-D5c, 1:100), MyHC type IIA (DSHB, SC-71c, 1:100), MyHC type IIB (DSHB, BF-F3c, 1:100), embryonic MyHC (eMHC; DSHB, F1.652s, 1:20), and laminin (Abcam, ab7463, 1:200). Primary antibodies were detected with appropriate Alexa Fluor–conjugated secondary antibodies (Invitrogen, 1:500 in PBS). Stitched mosaic brightfield and fluorescent images of entire cross-sections of the TA, SOL, and left ventricle of the heart were acquired using an automated microscope operating in upright configuration (Echo Revolution). TA and SOL myofiber morphology, centrally nucleated fiber counts, and fiber type composition was quantified using the MuscleJ 1.0.2 plugin in FIJI/ImageJ. Intramuscular immune cells including PMNs (Ly6G^+^ cells), total monocytes/MΦ (CD68^+^ cells), and M2-like MΦ (CD206^+^ cells) were quantified in cross-sections of the TA, SOL, and left ventricle of the heart either manually or using ImageJ (FIJI)-based automated detection with prominence thresholding^68^.

### RNA extraction, cDNA synthesis, and RT-qPCR

A portion of proximal half of snap frozen GAST muscle samples (∼50 mg) were homogenized in 1 mL of ice-cold TRIzol reagent (Invitrogen, 15596018) using a Fisher brand Bead Mill 4 Mini homogenizer (Thermo Fisher Scientific, 15-340-164) at 4 m/s for 45 s. Total RNA was extracted from muscle homogenates by phenol–chloroform phase separation followed by isopropanol precipitation, according to the manufacturer’s protocol with minor modifications. RNA concentration and purity were assessed by spectrophotometry using a NanoDrop 2000c UV-Vis Spectrophotometer (Thermo Scientific). To remove contaminating genomic DNA, RNA samples were treated with DNase I (Thermo Fisher Scientific, AM2222) followed by heat inactivation. One microgram of RNA was reverse transcribed to cDNA using the High-Capacity RNA-to-cDNA Kit (Applied Biosystems, 4387406). Reverse transcription-quantitative polymerase chain reaction (RT-qPCR) analysis was performed using a QuantStudio 5 Real-Time PCR System (Applied Biosystems, 384-well format). Reactions were carried out in 10 μL volumes using PowerUp™ SYBR™ Green Master Mix (Applied Biosystems, A25742) with gene-specific forward and reverse primers (1 μM each) and 1–40 ng of cDNA, depending on target gene abundance. All samples were analyzed in duplicate. Relative mRNA expression was calculated using the 2^-ΔΔCt^ method. Gene expression levels were normalized to 18S rRNA. Primer sequences used are listed in **Table 1**.

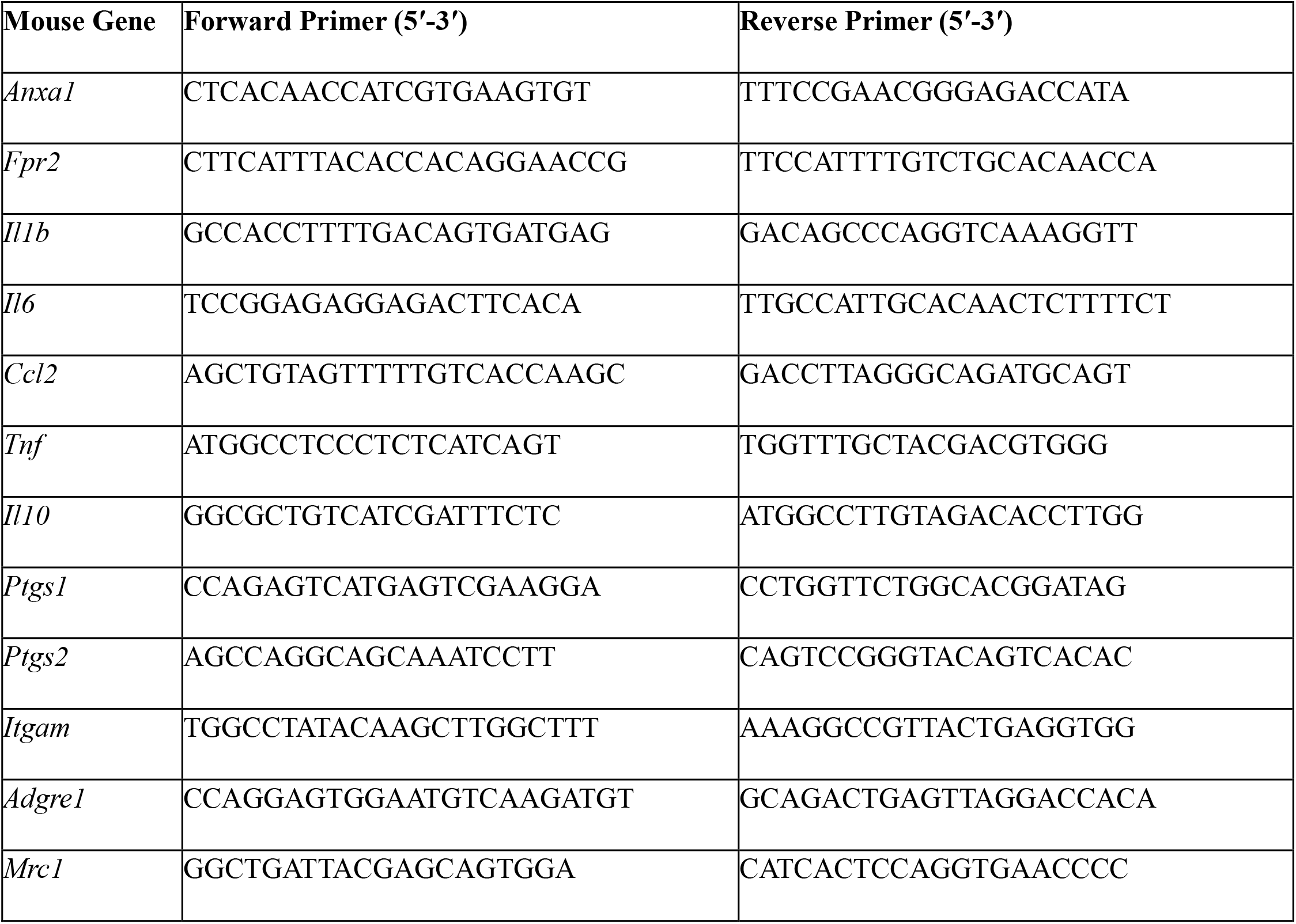
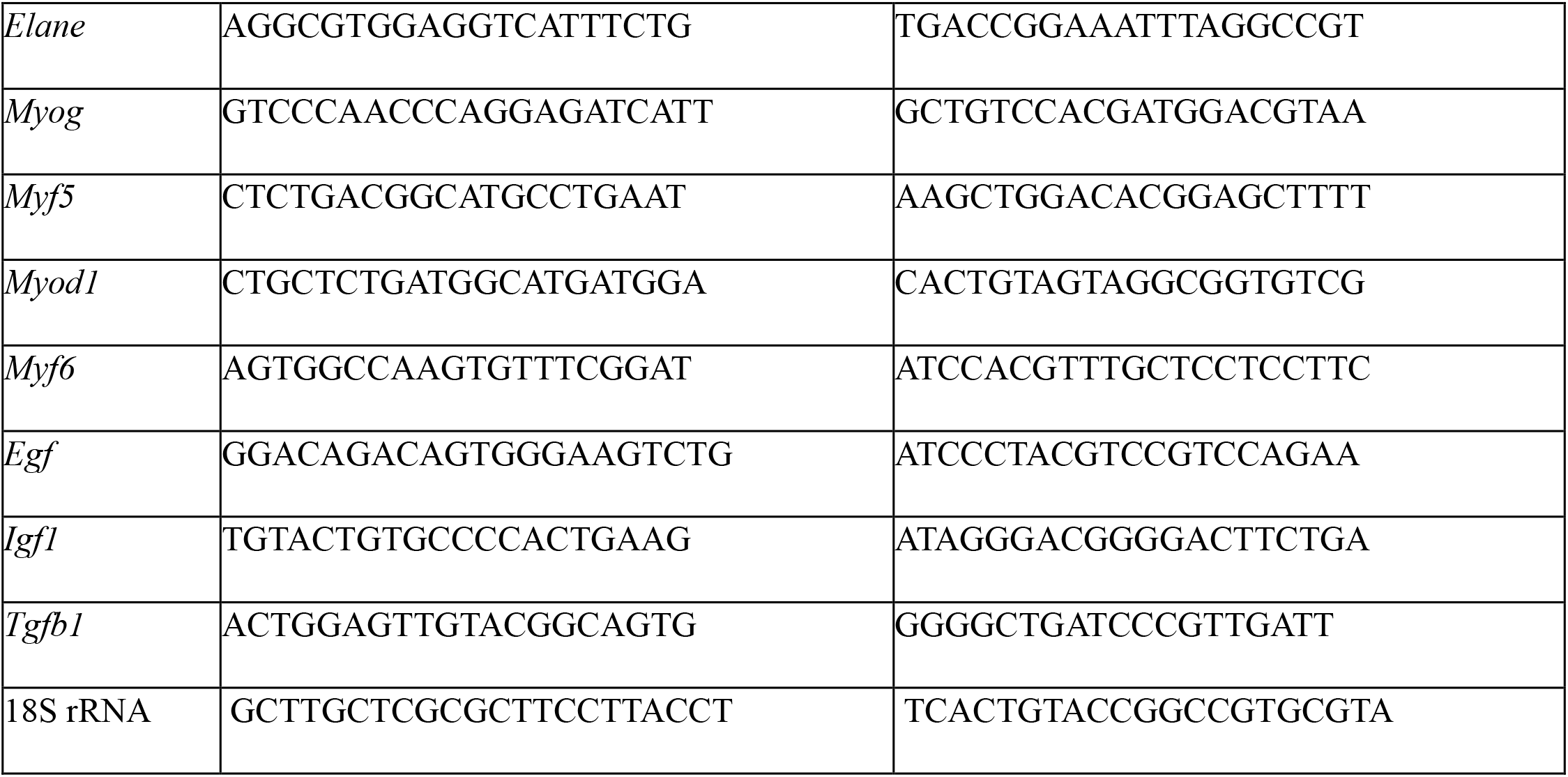

### Primary myoblast isolation and cell culture

Primary myoblasts were isolated from pooled hindlimb muscles of 6–8 week-old male D2.B10-Dmd*^mdx^*/J (D2-*mdx*) mice (*n* = 3) (Jackson Laboratories, Strain #:013141) using a previously described protocol (Hindi et al., 2017)^69^. Cells were cultured at 37°C in 5% CO₂ on T75 flasks coated with 10% Matrigel Matrix (Corning, 354234) in growth media (GM) consisting of Ham’s F-10 Nutrient Mixture (Gibco, 11550043) supplemented with 20% fetal bovine serum (FBS, Corning, 35-010-CV), basic fibroblast growth factor (bFGF, 10 ng/mL; PeproTech, 100-18B), and antibiotics [penicillin (100 U/mL) and streptomycin (100 µg/mL)]. Myoblasts (2 × 10⁵ cells/well) were seeded onto Matrigel-coated 12-well plates (Thermo Scientific, 130185) and maintained in GM for 72 hours to allow proliferation. Differentiation was then induced by switching to differentiation media (DM) consisting of Dulbecco’s modified Eagle medium (DMEM, Gibco, 11995-065) supplemented with 2% horse serum (HS; Gibco, 26050088) and antibiotics. Treatment groups included vehicle control (DMSO), deflazacort (10 µM), Ac2-26 (10 µM), or the FPR2 antagonist WRW4 (40 µM). Experimental treatments were prepared in DM, administered at the onset of myogenic differentiation, and maintained for 72 hours.

### Immunocytochemistry and image analysis

D2-*mdx* myotubes were fixed in 4% paraformaldehyde (PFA) at 4°C for 30 minutes, permeabilized with 0.1% Triton X-100 for 15 minutes at room temperature, and blocked in 1% bovine serum albumin (BSA) for 1 h at room temperature. Cells were then incubated overnight at 4°C with primary antibodies against sarcomeric myosin heavy chain (MyHC; DSHB, MF20c, 1:100) and myogenin (MyoG; DSHB, F5Dc, 1:100). The following day, cells were incubated with appropriate secondary antibodies, including Goat anti-mouse IgG2b Alexa Fluor 647 (Invitrogen, A-21242, 1:500) and Goat anti-mouse IgG1 Alexa Fluor 555 (Invitrogen, A-21127, 1:500), for 1 h at room temperature. Nuclei were counterstained with DAPI (Invitrogen, D21490, 2 μg/mL). Fluorescent images were acquired using an automated fluorescent microscope operating in inverted configuration (Echo Revolution). Images were captured from nine non-overlapping fields of view (FOVs) per well using a 10× Plan Fluorite objective. Images were analyzed using a custom ImageJ plugin developed in-house to quantify total cell density (DAPI^+^ nuclei/FOV), percentage of myogenin^+^ cells (% MyoG^+^/DAPI^+^ nuclei), and myotube area (% MyHC^+^ area/FOV).

## RESULTS

### Infiltrating myeloid cells are major sources of ANXA1 in dystrophic skeletal muscle

Tissue cross-sections of TA muscles from 8-week-old male *mdx* mice and age matched male wild type C57BL/6 mice were processed for immunohistochemistry staining to assess ANXA1 expression and cellular localization within dystrophic and healthy skeletal muscle, respectively (**Fig. 1**). Immunofluorescence analysis revealed robust ANXA1 staining of dystrophic muscle when compared to healthy muscle which showed relatively little ANXA1 expression (**Fig. 1A**). ANXA1 was found to be mainly co-localized with the large numbers of infiltrating myeloid cells present in dystrophic muscle. In inflamed regions lacking extensive MФ invasion or centrally located myonuclei, ANXA1 co-localized predominantly with PMNs (Ly6G^+^ cells) (**Fig. 1B**). In degenerative regions with extensive immune cell infiltration within the body of necrotic myofibers, ANXA1 expression was abundant and showed co-localization predominantly with monocytes/MФs (CD68^+^ cells) (**Fig. 1C**). In regenerative regions with an abundance of centrally nucleated myofibers and fewer PMNs or infiltrating MФs, ANXA1 expression was also detected predominantly in interstitial M2-like MФ (CD206^+^ cells) (**Fig. 1D**). Collectively, these findings demonstrate that infiltrating myeloid cell population, particularly PMNs and inflammatory monocytes/MФ, are major cellular sources of endogenous ANXA1 in dystrophic skeletal muscle. Notably, dystrophic myofibers themselves were generally not found to stain strongly for ANXA1 in the absence of inflammatory cell infiltration.

**Figure 1:**
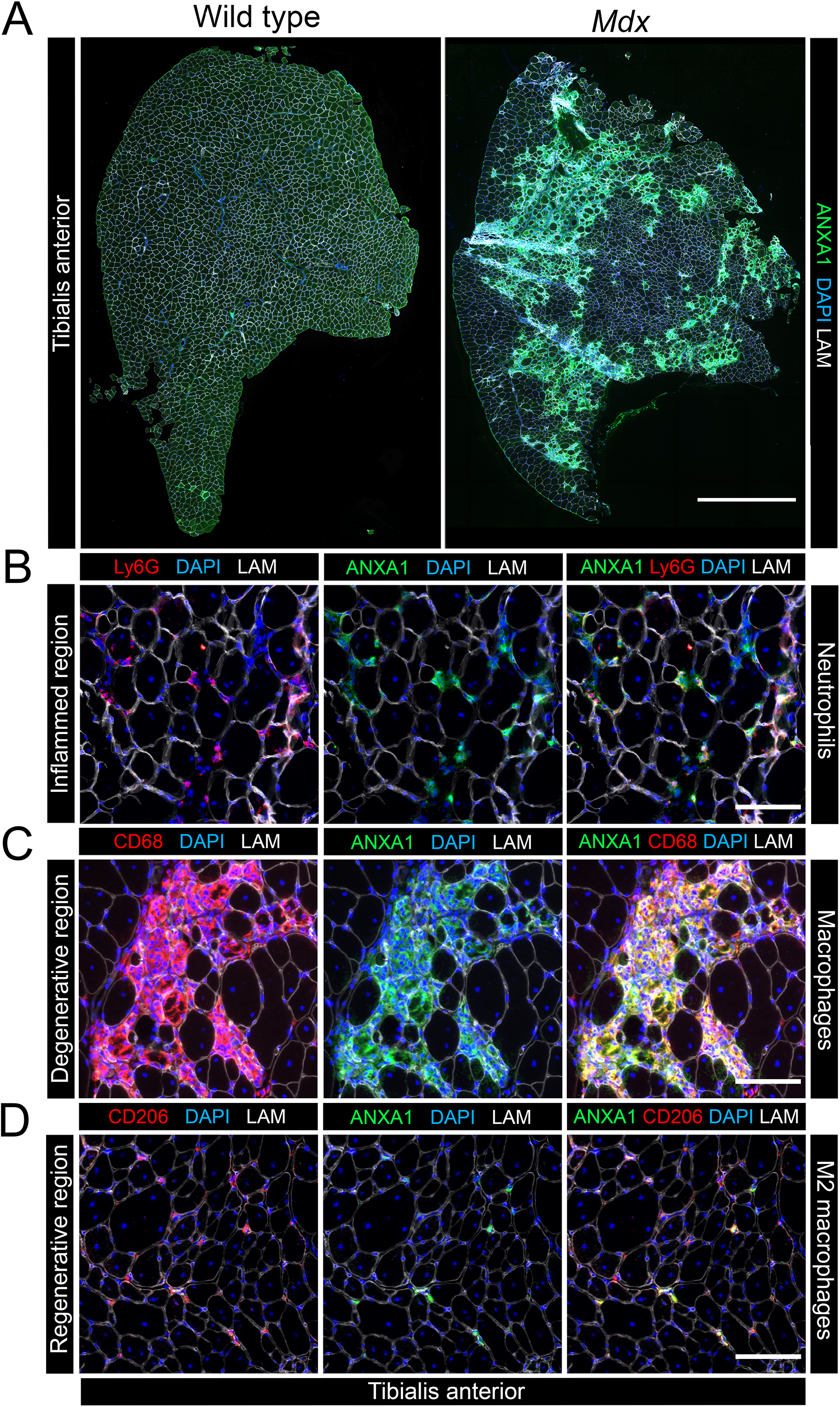
Co-localization of annexin A1 with infiltrating immune cells in dystrophic skeletal muscle. **A:** Representative stitched mosaic immunofluorescent images of whole tibialis anterior (TA) muscle cross-sections obtained from 8-week-old male wildtype (WT) and dystrophin-deficient *mdx* mice stained for annexin A1 (ANXA1; green), cell nuclei (DAPI; blue), and laminin (LAM; white). Scale bars on are 1000 µm. **B–D:** Higher-magnification images (20 ×) showing inflamed **(B)**, degenerative **(C)**, and regenerative **(D)** regions of interest within the *mdx* TA muscle. Specific cell types expressing high levels of ANXA1 (green) were identified as predominantly neutrophils (Ly6G^+^ cells; red) within the inflamed regions **(B)**, monocytes/macrophages (MΦ) (CD68^+^ cells; red) within the degenerative regions **(C)**, and M2-like MΦ (CD206^+^ cells; red) in regenerative regions **(D)**. Scale bars are 100 µm.

### Daily Ac2-26 treatment improves body composition and increases skeletal muscle mass in *mdx* mice

To test the effect of long-term systemic administration of ANXA1 mimetic peptides on dystrophic muscle pathology *in vivo* we treated 3-week-old *mdx* mice daily with vehicle control (0.25% DMSO), deflazacort (1 mg/kg/day), or Ac2-26 (1 mg/kg/day) for a duration of 5 weeks. Treatments were prepared in sterile saline and administered via daily IP injections. All mice gained body weight over time, but Ac2-26-treated mice showed significantly greater absolute body weights than mice treated with deflazacort between day 20 and 37 of the study (**Fig. S1A**). Analysis of the relative change in body weight over time revealed that deflazacort treated mice gained significantly less body weight than vehicle treated mice from day 9 to day 37 (**Fig. 2A**). In contrast, Ac2-26 treated mice gained significantly more body weight than vehicle treated mice from day 18 to 37 (**Fig. 2A**). Furthermore, Ac2-26 treated mice also gained significantly more body weight than deflazacort treated mice from day 7 to day 37 of the study (**Fig. 2A**). RT-qPCR analysis of the GAST muscle showed significantly higher expression of ANXA1 (*Anxa1*) mRNA in the deflazacort-treated group and a similar trend toward higher expression (*p* = 0.055) in the Ac2-26-treated group (**Fig. 2B**). Expression of the mRNA encoding the ANXA1 receptor FPR2 (*Frp2*) was readily detected in dystrophic skeletal muscle, but neither deflazacort nor Ac2-26 treatment influenced *Fpr2* expression (**Fig. 2C**). Terminal body weights were significantly greater in the Ac2-26 vs. deflazacort group, with a similar statistical trend observed for the Ac2-26 vs. vehicle groups (*p* = 0.056) (**Fig. 2D**). Terminal EchoMRI analysis of body composition at 8-weeks of age showed that the increased body weight in the Ac2-26 group was primarily attributable to accretion of lean mass, as Ac2-26-treated mice had significantly greater lean body mass than the deflazacort group and a similar trend was also observed when compared to the vehicle group (*p* = 0.055) (**Fig. 2E**). In contrast, absolute fat mass tended to be reduced in the Ac2-26 group when compared to vehicle group (*p* = 0.066) (**Fig. 2F**). Therefore, percent fat mass was also significantly lower in Ac2-26 vs. vehicle treated mice (**Fig. 2G**). In contrast, percentage lean mass did not differ between the three experimental groups (**Fig. 2H**).

**Figure 2:**
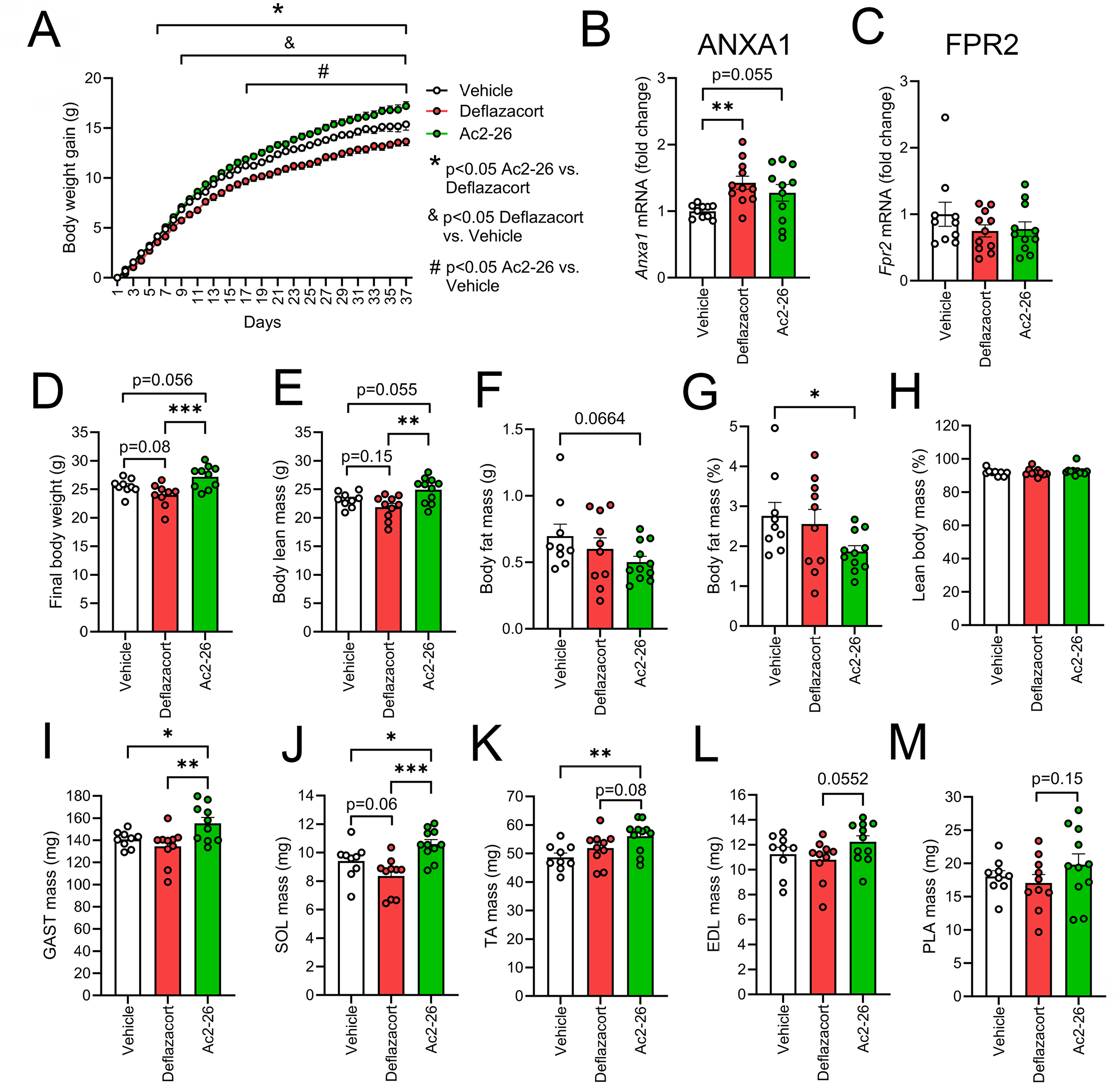
Effect of daily systemic treatment with deflazacort or Ac2-26 on body weight, body composition, and absolute muscle mass in *mdx* mice. **A:** Longitudinal tracking of cumulative body weight gain in male dystrophin deficient *mdx* mice that received once-daily intraperitoneal (IP) injections with vehicle control (0.25% DMSO), deflazacort (1 mg/kg/day), or Ac2-26 (1 mg/kg/day) from 3-weeks to 8-weeks of age. **B-C:** Endpoint reverse transcription-quantitative polymerase chain reaction (RT-qPCR) analysis of gastrocnemius (GAST) muscle mRNA expression of annexin A1 (ANXA1, *Anxa1*) **(B)** and formyl peptide receptor 2 (FPR2, *Fpr2*) **(C)**. **D-H:** Endpoint EchoMRI analysis of body composition including final body weight **(D),** absolute body lean mass **(E),** absolute body fat mass **(F),** percentage body fat mass **(G),** and percentage lean body mass **(H)**. **I-M:** Absolute wet weights of isolated individual hindlimb muscles including gastrocnemius (GAST) **(I),** soleus (SOL) **(J),** tibialis anterior (TA) **(K),** extensor digitorum longus (EDL) **(L)**, and plantaris (PLA**) (M)**. **A:** Data are presented as mean ± SEM (n = 10–11 mice per group). Statistical analysis was performed by two-way mixed model ANOVA followed by Fisher’s LSD post-hoc tests. \**p* < 0.05 for the Ac2-26 vs. deflazacort groups; & *p* < 0.05 for the deflazacort vs. vehicle groups; # *p* < 0.05 for the Ac2-26 vs. vehicle groups. **B-M:** Data are presented as the group mean ± SEM (n = 10–11 mice per group) with dot plots representing data from each individual mouse (biological replicates). Statistical analysis was performed by one-way ANOVA followed by Fisher’s LSD post-hoc tests. \**p* < 0.05, \*\**p* < 0.01, \*\*\**p* < 0.001 between the indicated experimental groups.

Consistent with these body composition findings, absolute terminal wet weights of major hindlimb muscles including the GAST (**Fig. 2I**), SOL (**Fig. 2J**), and TA (**Fig. 2K**) were significantly greater in Ac2-26-treated mice when compared to those receiving vehicle treatment. The mass of the GAST (**Fig. 2I**) and SOL (**Fig. 2J**) was also greater in the Ac2-26 vs. deflazacort groups, and similar statistical trends were observed for the TA (*p* = 0.08) (**Fig. 2K**), EDL (*p* = 0.052) (**Fig. 2L**), and PLA (*p* = 0.15) (**Fig. 2M**). When normalized to final body weights the relative mass of the TA muscle (mg/g body weight) was also greater in mice receiving either deflazacort or Ac2-26 when compared to the vehicle group (**Fig. S1B**). Moreover, relative SOL mass was greater in the Ac2-26 vs. deflazacort groups, with a similar trend observed for the Ac2-26 vs. vehicle groups (*p* = 0.068) (**Fig. S1C**). In contrast the relative masses of the GAST (**Fig. S1D**), PLA (**Fig. S1E**), and EDL (**Fig. S1F**) muscles did not differ between experimental groups.

### Daily Ac2-26 treatment improves the absolute strength of dystrophic skeletal muscle

Functional assessment of the *in vivo* strength of the ankle plantar flexors (GAST, PLA, and SOL) in live anesthetized mice at 8-weeks of age revealed that Ac2-26 treatment tended to improve maximal isometric twitch torque of the plantar flexor muscle group when compared to the deflazacort group (*p* = 0.053) (**Fig. 3A**). Analysis of the force-frequency relationship showed that Ac2-26-treated mice generated significantly greater isometric plantar flexion torque than deflazacort treated mice at stimulation frequencies between 80 and 300 Hz (**Fig. 3B**). Maximal isometric tetanic force of the ankle plantar flexors tended to be reduced when compared to the vehicle group in mice receiving daily deflazacort treatment (*p* = 0.06) and the Ac2-26 group was significantly stronger than the deflazacort group although there was no difference vs. vehicle control (**Fig. 3C**). We also performed terminal measurements of the *in situ* contractile properties of the TA muscle in live anaesthetized mice the day following *in vivo* functional testing prior to terminal tissue collection. Maximal isometric TA twitch force tended to be greater in the Ac2-26 vs. wither the deflazacort (*p* = 0.086) or vehicle (*p* = 0.10) groups (**Fig. 3D**). Analysis of the TA force-frequency relationship revealed that Ac2-26-treated mice generated greater maximal contractile force than vehicle-treated mice at stimulation frequencies between 150 and 300 Hz (**Fig. 3E**). Furthermore, the Ac2-26 group showed significantly greater absolute TA force generating capacity than the deflazacort group at stimulation frequencies between 80 and 300 Hz (**Fig. 3E**). Consistently, maximal isometric tetanic force of the TA muscle was significantly greater in the Ac2-26 group when compared to both the vehicle and deflazacort groups (**Fig. 3F**). To assess the effect of experimental treatments on the intrinsic contractile properties of the TA muscle, contractile force measurements were normalized to muscle physiological cross-sectional area (CSA) to calculate specific muscle force (mM/mm^2^). Maximal specific twitch (**Fig. 3G**), the specific force-frequency relationship (**Fig. 3H)**, and maximal specific tetanic force (**Fig. 3I**) did not differ significantly between experimental groups. Overall, these data show that long term systemic Ac2-26 treatment improves the absolute contractile function of dystrophic skeletal muscle predominantly via improvements in muscle mass and body composition.

**Figure 3:**
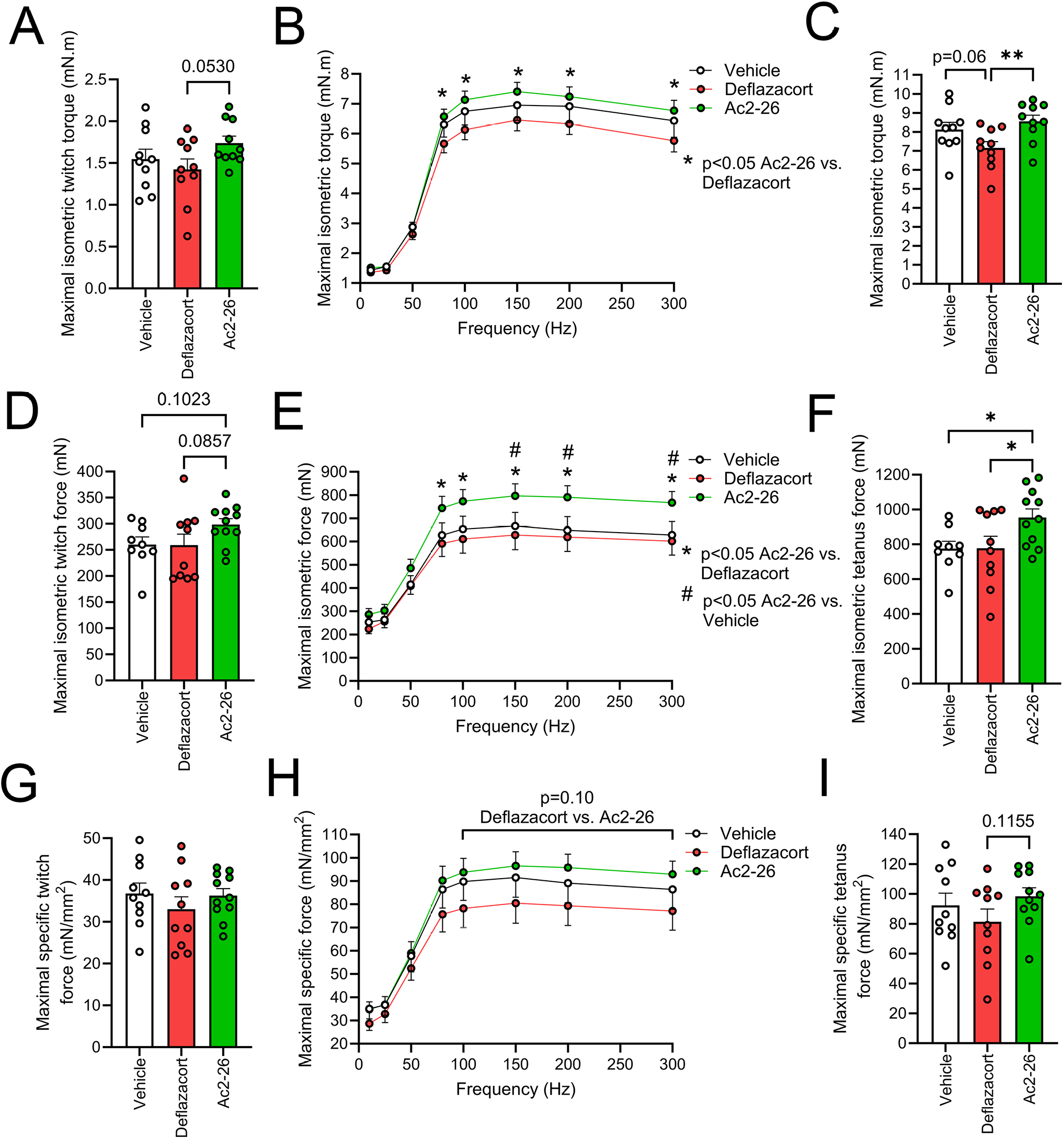
Effect of daily systemic treatment with deflazacort or Ac2-26 on contractile function of dystrophic skeletal muscle: **A–C:** *In vivo* ankle plantar flexion isometric torque (mN-m) parameters including maximal twitch **(A)**, the force-frequency relationship **(B),** and maximal tetanus **(C)** obtained from 8-week-old male dystrophin deficient *mdx* mice that received once-daily intraperitoneal (IP) injections with vehicle control (0.25% DMSO), deflazacort (1 mg/kg/day), or Ac2-26 (1 mg/kg/day) for a duration of 5-weeks. **D–F:** *In situ* absolute isometric force (mN) parameters of the tibialis anterior (TA) muscle including maximal twitch force **(D)**, the force-frequency relationship **(E)**, and maximal tetanic force **(F)**. **G–I:** *In situ* normalized TA specific force (mN/mm^2^) parameters including maximal specific twitch force **(G)**, specific force-frequency relationship **(H)**, and maximal specific tetanic force **(I)**. **A, C, D, F, G, I:** Data are presented as the mean ± SEM (n = 10–11 mice per group) with dot plots representing data from each individual mouse (biological replicates). Statistical analysis was performed using a one-way ANOVA followed by Fisher’s LSD post-hoc test. *p < 0.05, **p < 0.01. **B, E, H:** Data are presented as the group mean ± SEM (n = 10–11 mice per group). Statistical analysis was performed by two-way mixed model ANOVA followed by Fisher’s LSD post-hoc test. \**p* < 0.05 for Ac2-26 vs. deflazacort groups; # *p* < 0.05 for Ac2-26 vs. vehicle groups.

### Effect of intermittent deflazacort and Ac2-26 treatment in *mdx* mice

Recent studies have suggested that intermittent glucocorticoid therapy may benefit dystrophic muscle pathology while having less negative side effects when compared to more than traditional daily treatment regimens^70–73^. Therefore, we also compared the effect of once-weekly IP injections of 3-week-old *mdx* with vehicle control (0.25% DMSO), deflazacort (1 mg/kg/week) or Ac2-26 (1 mg/kg/week) for a duration of 5 weeks. Unlike our daily treatment regimen experiments, absolute body weight (**Fig. S2A**) and relative gain in body weight (**Fig. S2B**) did not significantly differ between experimental groups at any time-point. Nevertheless, final terminal body weight did still tend to be reduced in the deflazacort group when compared to the vehicle group (*p* = 0.08) and was significantly greater in mice receiving weekly treatment with Ac2-26 when compared to those receiving deflazacort (**Fig. S2C**). There were no significant effects of weekly deflazacort or Ac2-26 treatments on absolute lean mass (**Fig. S2D**) or percentage of lean mass (**Fig. S2E**). Nevertheless, absolute fat mass tended to be lower in the Ac2-26 group than the deflazacort group (*p* = 0.09) (**Fig. S2F**). Thus, mice receiving once-weekly Ac2-26 treatment showed significantly lower percentage fat mass than those receiving weekly deflazacort (**Fig. S2G**). Despite the lack of effect of weekly treatments on overall lean body mass, mice receiving once-weekly treatment with Ac2-26 did still show greater absolute mass of the TA (**Fig. S2H**) and SOL (**Fig. S2I**) when compared to those receiving one-weekly deflazacort. Similar statistical trends were also observed for the PLA (p= 0.06) (**Fig. S2J**) and EDL (p=0.11) (**Fig. S2K**). Once-weekly treatment with deflazacort, but not Ac2-26, also tended to reduce GAST mass (*p* = 0.05), a negative effect not observed in response to once-weekly Ac2-26 administration (**Fig. S2L**). When muscle weights were normalized to body weight, the mice receiving once-weekly treatments with Ac2-26 group tended to have greater relative TA mass than the vehicle group (*p* = 0.084) (**Fig. S2M**) and showed significantly greater relative SOL mass (**Fig. S2N**) when compared to the deflazacort group. However, this was not observed for the PLA (**Fig. S2O**) or EDL (**Fig. S2P**). Finally, relative GAST mass was greater than vehicle control in mice receiving once-weekly treatment with Ac2-26, but not an equivalent once-weekly dose of deflazacort (**Fig. S2Q**).

Once-weekly treatment with deflazacort or Ac2-26 had minimal apparent effect of *in vivo* plantar flexion torque parameters (**Fig. S3A-C**). Nevertheless, once-weekly deflazacort and Ac2-26 treatment tended to have overall opposing effects on maximal *in situ* TA twitch force with deflazacort treated mice tending to have lower twitch force than those receiving either vehicle (*p* = 0.063) or Ac2-26 (*p* = 0.058) (**Fig. S3D**). Consistently, once-weekly deflazacort treatment reduced the *in situ* contractile force of the TA muscle at stimulation frequencies between 80 and 300 Hz when compared to either the vehicle or Ac2-26 groups (**Fig. S3E**). Mice receiving once-weekly deflazacort treatment also displayed significantly lower maximal isometric tetanic force than either the vehicle or Ac2-26 groups, although those receiving Ac2-26 were not significantly stronger than those receiving vehicle control (**Fig. S3F**). Normalization of absolute TA contractile force measurements to pCSA reduced the magnitude and statistical significance of the differences between groups for TA specific contractile force (**Fig. S3G-H**). Nevertheless, mice receiving once-weekly Ac2-26 treatment did still tend to show greater maximal specific isometric force than those receiving once-weekly deflazacort treatment (*p* = 0.053) (**Fig. S3I**). Overall, these data show that consistent with prior reports, that once-weekly glucocorticoid treatment regimens appear to have overall less deleterious effect on body weight gain in *mdx* mice than daily corticosteroid administration regimens. Nevertheless, once-weekly Ac2-26 still elicited superior beneficial effects on body composition, muscle mass, and absolute TA contractile strength when compared to once-weekly treatment with an equivalent dose of deflazacort. While still apparently quite effective, once-weekly Ac2-26 treatment clearly did not have as obvious benefits as daily treatment in *mdx* mice. Therefore, we focused further histological and molecular analysis on muscle tissue samples obtained from *mdx* mice receiving daily treatment regimens.

### Effects of daily deflazacort and Ac2-26 treatment on dystrophic skeletal muscle histopathology in *mdx* mice

To assess the effect of Ac2-26 on the gross histology of dystrophic skeletal muscle, tissue cross-sections of fast-twitch TA muscles from 8-week-old *mdx* mice that had received once-daily treatment with deflazacort or Ac2-26 for a duration of 5 weeks were stained with H & E and imaged via brightfield microscopy (**Fig. 4A**). The percentage of regenerating (centrally nucleated) myofibers was significantly greater in muscle cross-sections obtained from mice receiving Ac2-26 treatment when compared to vehicle and deflazacort treated mice (**Fig. 4B**). The proportion of total myofibers containing one single centrally located nucleus did not differ between experimental groups (**Fig. 4C**). Nevertheless, the proportion of total myofibers containing two central nuclei (**Fig. 4D**) and ≥3 central nuclei (**Fig. 4E**) were both significantly greater in the Ac2-26 group when compared to mice receiving either vehicle or deflazacort treatment. Analysis of the frequency distribution of myofiber cross-sectional area (CSA) revealed that both deflazacort and Ac2-26 treatments reduced the percentage of myofibers with a CSA between 1000-2000 µm^2^ (**Fig. 4F**). However, only Ac2-26 significantly increased the proportion of myofibers with a CSA between 3000-4000 µm^2^ (**Fig. 4F**). We also analyzed muscle fiber type profile and myofiber type-specific CSA in TA muscle cross-sections obtained from *mdx* mice treated daily with vehicle, deflazacort, or Ac2-26 (**Fig. 4G**). The total number of myofibers present within TA muscles did not differ between experimental groups (**Fig. 4H**). However, the mean myofiber CSA of *mdx* mice receiving either deflazacort or Ac2-26 were significantly larger than the vehicle control group (**Fig. 4I**). Treatment with deflazacort or Ac2-26 did not influence the relative proportion of various muscle fiber types within the TA muscle (**Fig. 4J**). Nevertheless, mice treated with either deflazacort or Ac2-26 displayed significantly greater average CSA of the type IIA, IIX, and IIB myofiber populations within the TA muscle (**Fig. 4K**). In contrast, the average CSA of the type I myofiber population did not differ between groups (**Fig. 4K**).

**Figure 4:**
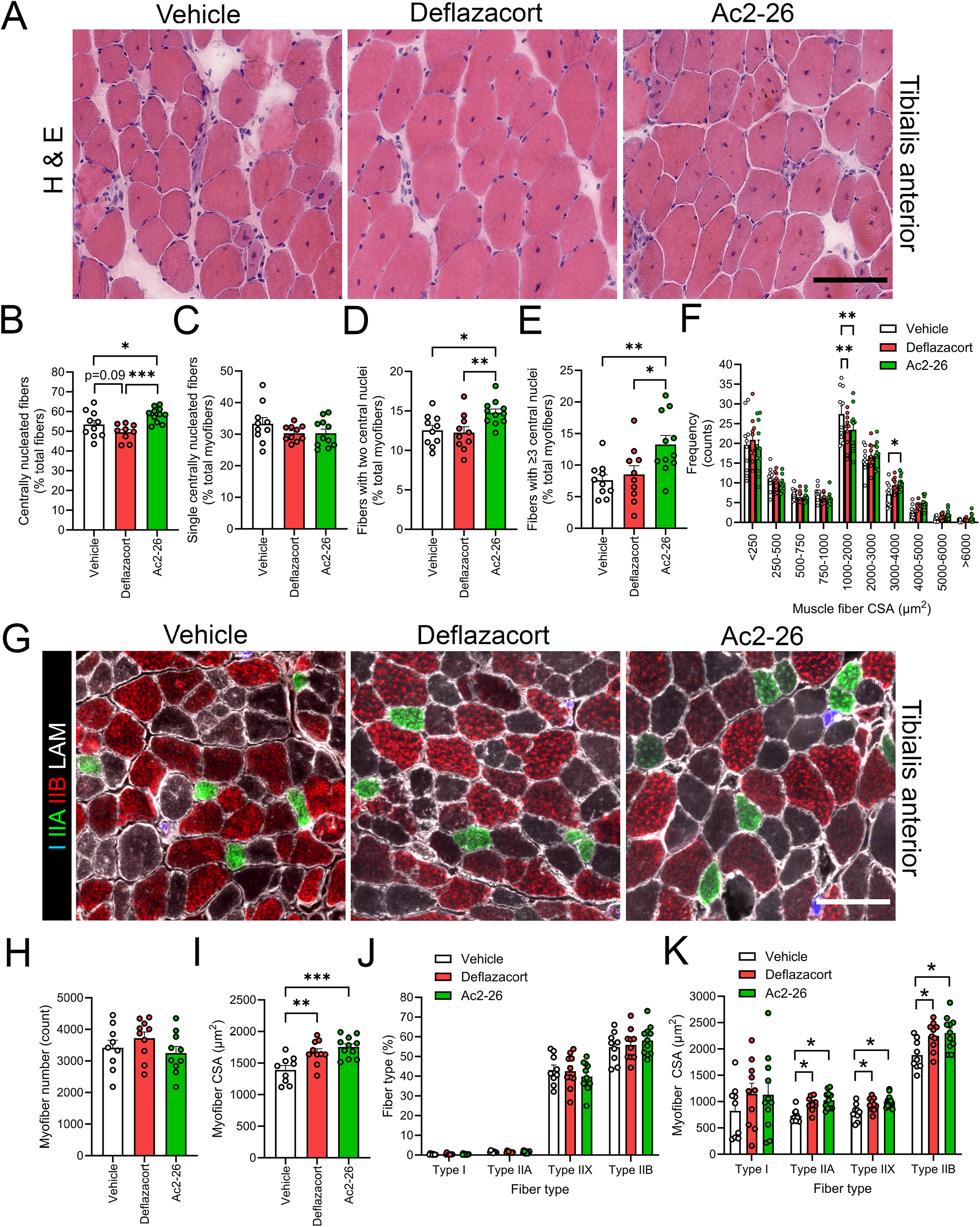
Histological and immunofluorescence analysis of fast-twitch tibialis anterior myofiber regeneration and muscle fiber type composition of *mdx* mice receiving daily systemic treatment with deflazacort or Ac2-26. **A:** Representative hematoxylin and eosin (H & E) stained cross-sections of tibialis anterior (TA) muscles obtained from 8-week-old male dystrophin deficient *mdx* mice that received once-daily intraperitoneal (IP) injections with vehicle control (0.25% DMSO), deflazacort (1 mg/kg/day) or Ac2-26 (1 mg/kg/day) for a duration of 5-weeks. **B–F:** Quantification of the total proportion of regenerating (centrally nucleated) myofibers **(B)**, percentage of single centrally nucleated fibers **(C)**, percentage of fibers with two central nuclei **(D)**, percentage of fibers with ≥3 central nuclei **(E)**, and frequency distribution of muscle fiber cross-sectional area (CSA) **(F)**. **G:** Representative immunofluorescence images of TA muscle cross-sections stained with primary antibodies against for type I myosin heavy chain (MyHC; blue), type IIA MyHC (green), type IIB (red), and laminin (LAM; white). Type IIX fibers (black) are identified by lack of positive staining for type I, IIA, or IIB MyHC. **H–K:** Quantitative analysis of total myofiber number **(H)**, overall mean myofiber CSA **(I)**, fiber type percentage distribution **(J)**, and fiber type-specific myofiber CSA **(K)**. Data is presented as the group mean ± SEM (n = 10–11 mice per group) with dot plots representing data from each individual mouse (biological replicates). Statistical analysis was performed by one-way ANOVA followed by Fisher’s LSD post-hoc tests. \**p* < 0.05, \*\**p* < 0.01, \*\*\**p* < 0.001 between the indicated experimental groups. Scale bars are 100 µm.

We also assessed the effect of daily deflazacort vs. Ac2-26 treatment on the histology of the predominantly slow-twitch SOL muscle (**Fig. S4A**). The overall percentage of regenerating (centrally nucleated) myofibers (**Fig. S4B**) and proportion of regenerating myofibers containing a single centrally located nucleus (**Fig. S4C**) did not differ between experimental groups. Nevertheless, the proportion of total myofibers with two centrally located nuclei was significantly increased in mice receiving Ac2-26 treatment when compared to vehicle control and a similar statistical trend observed between the Ac2-26 vs. deflazacort groups (*p* = 0.08) (**Fig. S4D**). Daily treatment with deflazacort, but not Ac2-26, had overall deleterious effect on the frequency of distribution of SOL myofiber CSA (**Fig. S4E**). Specifically, deflazacort treatment increased the proportion of smaller myofibers in the CSA range of 500-1000 µm^2^ while decreasing the proportion of large myofibers with a CSA between 2000-3000 µm^2^ (**Fig. S4E**). We also assessed the effect of daily deflazacort and Ac2-26 treatment on SOL muscle fiber type in *mdx* mice (**Fig. S4F)**. There were no significant differences in total SOL muscle fiber number (**Fig. S4G**). However, mean myofiber CSA was significantly increased in mice receiving Ac2-26 treatment when compared to either the vehicle or deflazacort groups (**Fig. S4H**). Treatment with deflazacort or Ac2-26 both resulted in a slow-to-fast fiber type switch as indicated by a decreased percentage of type IIA and increased percentage of IIX myofibers in the SOL, respectively (**Fig. S4I**). However, only mice receiving Ac2-26 treatment showed an increase in the mean CSA of the type I, IIA, and IIX myofiber populations when compared to those receiving vehicle or deflazacort (trend *p* = 0.07 for type I fibers vs. deflazacort) (**Fig. S4J**). In contrast, SOL type IIB myofiber CSA did not differ between experimental groups (**Fig. S4J**). Overall, these data show that daily Ac2-26 treatment appears to have relatively greater beneficial effects than deflazacort in the predominantly slow-twitch SOL muscle.

### Daily deflazacort and Ac2-26 treatment have equivalent suppressive effects on inflammation of dystrophic muscle

One of the primary mechanisms that has been proposed to mediate the benefits of glucocorticoid treatment in DMD is their potent anti-inflammatory effects upon the various immune cell populations that accumulate within dystrophic muscle tissue^74^. Therefore, we compared the effect of daily administration of deflazacort and Ac2-26 on inflammation of the fast-twitch TA muscle in *mdx* mice (**Fig 5A**). Mice treated once-daily with either deflazacort or Ac-2-26 showed a marked reduction in the number of PMNs (Ly6G^+^ cells) within TA muscle cross-sections (**Fig. 5B**). Total TA numbers of monocytes/MФ (CD68^+^ cells) were also significantly reduced in response to treatment with either deflazacort or Ac2-26 (**Fig. 5C**). In contrast, absolute numbers of TA M2-like MФ (CD206^+^ cells) did not differ between experimental groups (**data not shown**). Therefore, the relative number of M2-like MФ (CD206^+^ cells) as a proportion of total monocytes/MФ (CD68^+^ cells) within the TA was significantly increased in *mdx* mice receiving deflazacort or Ac2-26 treatment (**Fig. 5D**). There were no significant differences between the deflazacort and Ac2-26 groups for intramuscular infiltration of the TA muscle by PMNs (**Fig. 5B**), total monocyte/MФ (**Fig. 5C**), M2-like MФ (**data not shown**), or the percentage of M2-like MФ (**Fig. 5D**).

**Figure 5:**
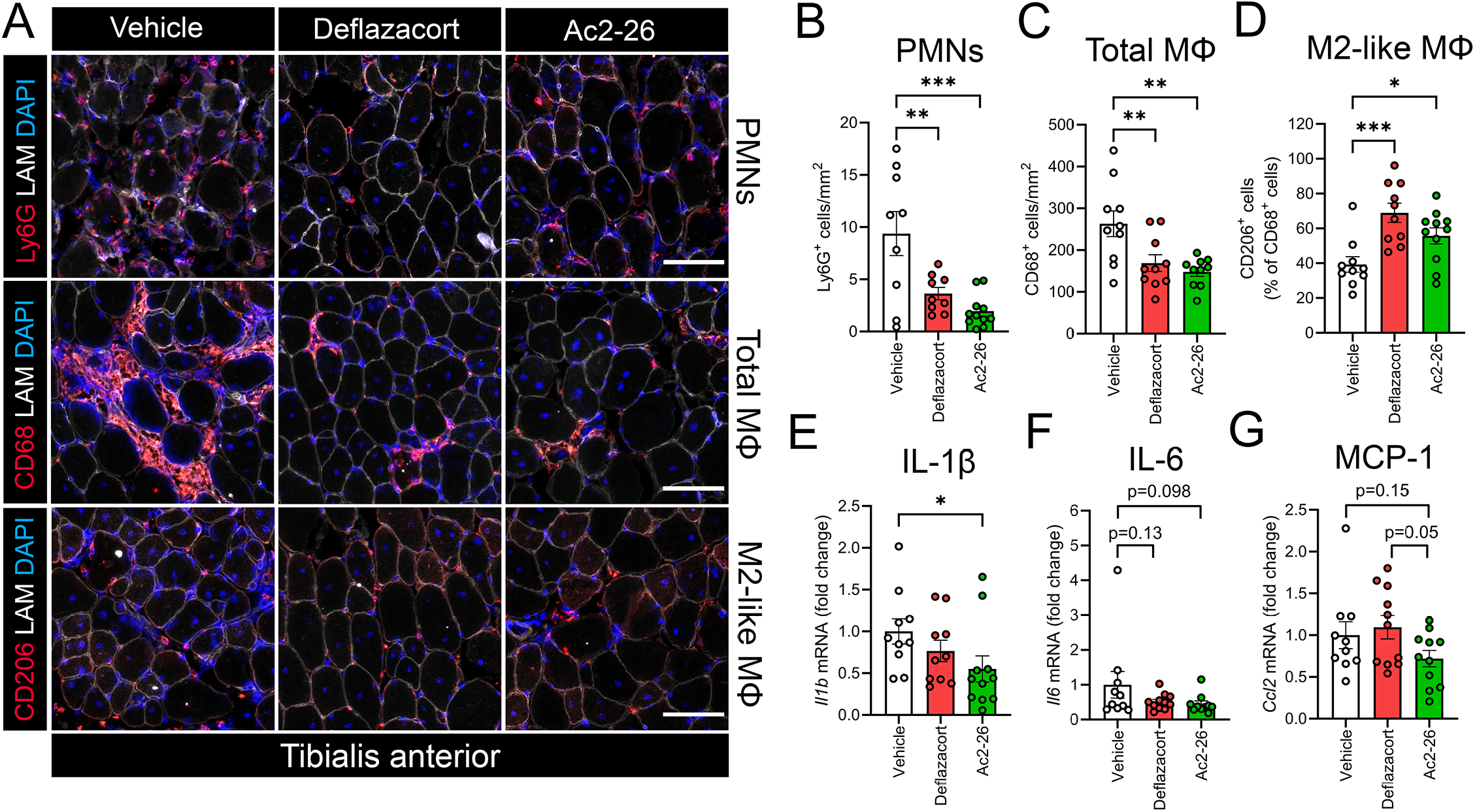
Immunofluorescence analysis of infiltrating immune cell populations in fast-twitch tibialis anterior muscles of *mdx* mice following daily systemic deflazacort or Ac2-26 administration. **A:** Representative immunofluorescent images of tibialis anterior (TA) muscle cross-sections from 8-week-old male dystrophin-deficient *mdx* mice that had received daily intraperitoneal (IP) injections with vehicle control (0.25% DMSO), deflazacort (1 mg/kg/day, or Ac2-26 (1 mg/kg/day) for a duration of 5-weeks were stained for distinct immune cell markers (red), laminin (LAM; white), and cell nuclei (DAPI; blue). Rows from top to bottom show neutrophils (Ly6G^+^ cells; red), total monocytes/macrophages (MΦ) (CD68^+^ cells; red), and M2-like MΦ (CD206^+^ cells; red) across the treatment groups. **B–D:** Quantitative analysis of intramuscular numbers of neutrophils (Ly6G^+^ cells) **(B)**, total monocytes/ MΦ (CD68^+^ cells) **(C)**, and the proportion of M2-like MΦ relative to total monocytes/MΦ (CD68^+^ cells) **(D)**. **E-F:** Reverse transcription-quantitative polymerase chain reaction (RT-qPCR) analysis of gastrocnemius (GAST) muscle mRNA expression of interleukin-1 beta (IL-1β, *Il1b*) **(E),** interleukin-6 (IL-6, *Il6*) **(F)**, and monocyte chemoattractant protein 1 (MCP-1, *Ccl2*). Data are presented as the group mean ± SEM (n = 10–11 mice per group). Statistical analysis was performed using a one-way ANOVA followed by Fisher’s LSD post-hoc test. \**p* < 0.05, \*\**p* < 0.01, \*\*\**p* < 0.001 between the indicated experimental groups. Scale bars are 100 µm.

RT-qPCR analysis of the predominantly fast-twitch GAST muscle showed that daily treatment with Ac2-26 significantly reduced mRNA expression of the major pro-inflammatory cytokine interleukin 1 beta (IL-1β, *Il1b)* when compared to the vehicle group (**Fig. 5E**). A similar statistical trend (*p* = 0.098) was also observed towards a reduction in interleukin-6 (Il-6, *Il6*) mRNA expression in mice receiving Ac2-26 treatment (**Fig. 5F**). Neither deflazacort nor Ac2-26 influenced the expression of the pro-inflammatory cytokine tumor necrosis factor alpha (TNF-α, *Tnf*) (**Fig. S5A**), the anti-inflammatory cytokine interleukin-10 (IL-10, *Il10*) (**Fig. S5B**), or the pro-inflammatory lipid mediator biosynthetic enzyme cyclooxygenase-2 (COX-2, *Ptgs2*) **(Fig. S5C)**. Nevertheless, daily treatment with Ac2-26 did increase mRNA expression of constitutive lipid mediator biosynthetic enzyme cyclooxygenase 1 (COX-1, *Ptgs1*) relative to the vehicle group (**Fig. S5D**). Among immune cell marker genes, mRNA expression of the pan innate immune cell marker CD11b (*Itgam*) did not differ between experimental groups **(Fig. S5E)**. However, expression of the mature MФ marker F4/80 (*Adgre1*) was increased in mice receiving Ac2-26 compared to both the vehicle and deflazacort groups **(Fig. S5F)**. In contrast, expression of the chemokine monocyte chemoattractant protein 1 (MCP-1, *Ccl2*) **(Fig. 5G)** and the M2-like MФ marker CD206 (*Mrc1*) **(Fig. S5G)** was reduced in response to Ac2-26 treatment when compared to the deflazacort group. Expression of the PMN marker neutrophil elastase (ELANE, *Elane*) was also reduced in mice receiving deflazacort compared to both the vehicle and Ac2-26 groups **(Fig. S5H)**. Among myogenic regulatory factor genes, deflazacort treatment increased expression of myogenin (*Myog*) (**Fig. S5I**), whereas neither deflazacort nor Ac2-26 influenced myogenic factor 5 (Myf-5, *Myf5*) (**Fig S5J**), myoblast determination protein 1 (MyoD1, *Myod1*) (**Fig, S5K**), or myogenic regulatory factor 4 (MRF4, *Myf6*) **(Fig. S5L)**. Finally, deflazacort treatment increased mRNA expression of major growth factor genes including epidermal growth factor (EGF, *Egf*) (**Fig. S5M**) and insulin-like growth factor 1 (IGF-1, *Igf1*) (**Fig. S5N**). Nevertheless, expression of transforming growth factor-beta (TGF-β, *Tgfb1*) did not differ between experimental groups (**Fig. S5O**).

We also assessed the effect of deflazacort and Ac2-26 treatment on innate immune cell infiltration of the predominantly slow-twitch SOL muscle (**Fig. S6A**). Both deflazacort and Ac2-26 treatment significantly reduced PMN (Ly6G^+^ cell) counts in the SOL muscle (**Fig. S6B**). However, neither deflazacort nor Ac2-26 influenced the total number of monocytes/MФ (CD68^+^ cells) within the SOL muscle (**Fig. S6C**). Furthermore, daily treatment with Ac2-26 significantly reduced M2-like MФ (CD206^+^ cell) number in the SOL when compared to both the vehicle and deflazacort group (**Fig. S6D**).

A recent study showed that the once-daily treatment of juvenile D2-*mdx* mice with the small molecule FRP2 agonist BMS-986235 was able to stimulate resolution of the dystrophic heart^75^. Therefore, we also tested the effect of daily deflazacort or Ac2-26 treatment on immune cell infiltration of dystrophic cardiac muscle in our B10-*mdx* mice (**Fig. S7A**). Once-daily treatment with either deflazacort or Ac2-26 reduced PMN (Ly6G^+^ cell) numbers in the left ventricle of the heart (**Fig. S7B**). However, neither deflazacort or Ac2-26 significantly influenced total numbers of monocytes/MФ (CD68^+^ cells) (**Fig. S7C**), or the number of M2-like MФ (CD206^+^ cells) (**data not shown**). Nevertheless, the relative number of M2-like MФ (CD206^+^ cells) as a proportion of total monocytes/MФ (CD68^+^ cells) tended to be increased in hearts of mice receiving Ac2-26 treatment relative to the vehicle group (**Fig. S7D**). Overall, these data show that Ac2-26 mimics the anti-inflammatory effects of deflazacort upon dystrophic muscle tissue. However, the effect of both deflazacort and Ac2-26 to reduce monocyte/MФ infiltration appears to be specific to the fast-twitch TA muscle.

### Ac2-26 directly stimulates *in vitro* myogenesis of dystrophic skeletal myoblasts

To elucidate the potential cell types that might mediate the beneficial effects of Ac2-26 treatment on dystrophic skeletal muscle pathology we isolated primary myoblasts from pooled hindlimb muscles obtained from dystrophin-deficient D2-*mdx* mice. To test the potential role of endogenous extracellular ANXA1 signaling we first treated dystrophic myoblasts during 72 h of myogenic differentiation with WRW4 (40 µM), a specific pharmacological inhibitor of the ANXA1 extracellular receptor FPR2, or its corresponding vehicle control (0.4% DMSO) (**Fig. 6A**). WRW4 treatment reduced overall primary skeletal muscle cell density (**Fig. 6B**) and the total number of differentiated myogenic cells (MyoG^+^/mm^2^) (**Fig. 6C**) but did not affect the relative percentage of differentiation (% MyoG^+^ cells) due to the simultaneous decrease in total cell number (**Fig. 6D**). Furthermore, WRW4 treatment had an overall marked deleterious effect on myotube formation as evidenced by a significant reduction in myotube (% MyHC^+^ cell) area per field of view (FOV) following 72 h of myogenic differentiation (**Fig. 6E**). Overall, these data show that endogenous FPR2-dependent signaling is essential for the normal myogenic differentiation in dystrophic skeletal muscle cells.

**Figure 6:**
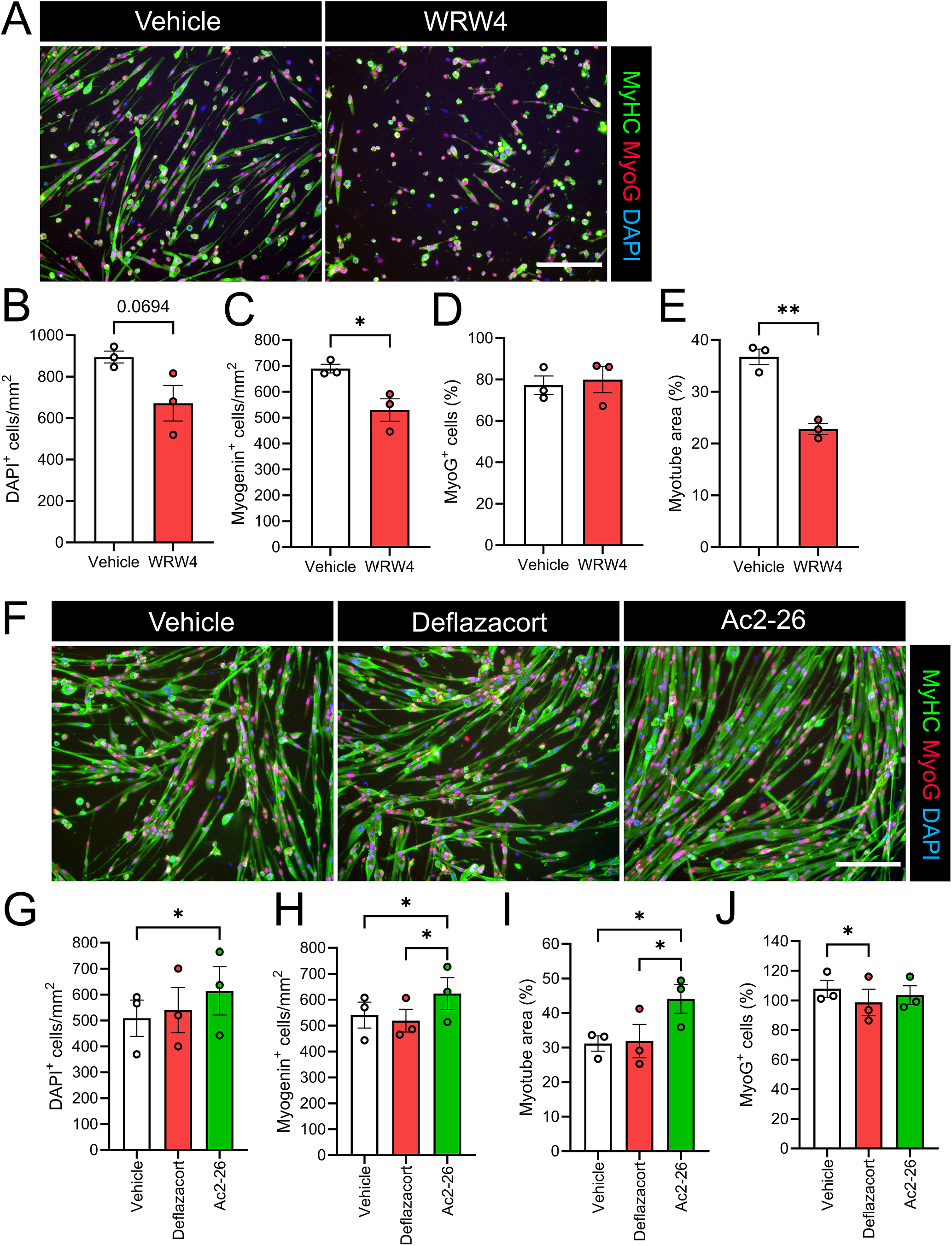
Direct role of formyl peptide receptor 2 signaling in *in vitro* myogenesis of dystrophic skeletal myoblasts. **A:** Representative immunofluorescence images of differentiating D2-*mdx* primary myoblast cultures treated for 72 h with vehicle control (0.4% DMSO) or the pharmacological formyl peptide receptor 2 (FPR2) antagonist WRW4 (40 µM). The resulting myotube cultures were fixed in 4% paraformaldehyde (PFA) and stained with primary antibodies against sarcomeric myosin heavy chain (MyHC; green) and myogenin (MyoG; red). Cell nuclei were counterstained with DAPI (blue). Quantitative image analysis of total (DAPI^+^) cell density **(B)**, total differentiated myoblasts (myogenin^+^ cells) **(C)**, the percentage of total cells expressing myogenin (% MyoG^+^ cells) **(D)**, and percentage of myotube (myosin^+^) cell area per field of view (FOV) (**E**). **F:** Representative immunofluorescence images of differentiating primary D2-*mdx* myoblast cultures treated for 72 h with vehicle control (0.02% DMSO), deflazacort (10 µM), or Ac2-26 (10 µM). The resulting myotube cultures were fixed in 4% PFA and stained with primary for sarcomeric MyHC (green), MyoG (red), and DAPI (blue). Quantitative image analysis of total (DAPI^+^) cell density **(G)** differentiated (MyoG^+^) cell density **(H)**, percentage myotube area per FOV **(I)**, and percentage of differentiated (% MyoG^+^) cells **(J)**. Data are presented as mean ± SEM of primary myoblasts isolated from n=3 separate mice with dots showing each data obtained from each individual biological replicate (mouse). **B-E:** Statistical analysis was performed by two-tailed unpaired t-tests. **G-J:** Statistical analysis was performed by one-way ANOVA followed by Fisher’s LSD post-hoc tests. \**p* < 0.05, \*\**p* < 0.01 between the indicated experimental groups. Scale bars are 200 µm.

We next tested the effect of treatment of differentiating D2-*mdx* myoblasts with an equivalent dose of deflazacort (10 µM), Ac2-26 (10 µM), or matching vehicle control (0.02% DMSO) (**Fig. 6F**). Following 72 h of myogenic differentiation, Ac2-26 treatment increased overall cell density (**Fig 6G**), the absolute number of differentiated (MyoG^+^/mm^2^) myogenic cells (**Fig. 6H**), and the overall extent of myotube formation (% MyHC^+^ cell area/FOV) (**Fig. 6I**) when compared to cells receiving vehicle control treatment. Ac2-26 treatment also increased the absolute amount of myogenic differentiation (MyoG^+^/mm^2^) (**Fig. 6H**) and overall myotube formation (% MyHC^+^ cell area/FOV) when compared to deflazacort treated cells, which themselves did not differ significantly from vehicle control. Finally, treatment with deflazacort, but not Ac2-26, reduced the relative proportion of cells undergoing myogenic differentiation (% MyoG^+^ cells) on day 3 of differentiation (**Fig. 6J**). Overall, these data show that the ANXA1 mimetic peptide Ac2-26 has more potent direct pro-myogenic effects on dystrophic myoblasts when compared to deflazacort treatment.

## DISCUSSION

In the current study, we investigated the effect of 5 weeks of systemic treatment with the ANXA1 mimetic peptide Ac2-26 on dystrophic skeletal muscle pathology in the preclinical *mdx* mouse model of DMD. We further compared the effects of Ac2-26 with the synthetic glucocorticoid deflazacort as the current standard of care for human DMD patients. Endogenous ANXA1 was found to be localized predominantly to infiltrating myeloid cells in dystrophic muscle. Daily systemic Ac2-26 treatment reproduced the anti-inflammatory effects of deflazacort while providing broader benefits in terms of body composition, skeletal muscle mass, myofiber size, and absolute muscle strength. Ac2-26 treatment also directly stimulated myogenesis in primary *D2-mdx* myoblasts, whereas pharmacological inhibition of endogenous FPR2 signaling impaired basal dystrophic myotube formation. Overall, these findings show that the ANXA1–FPR2 pathway is an important regulator of both inflammation-resolution and myofiber regeneration within dystrophic skeletal muscle and suggest that ANXA1 mimetic peptides such as Ac2-26 may provide novel steroid-sparing therapeutics for human DMD patients.

ANXA1 has been previously shown to be highly expressed by the various infiltrating myeloid cell populations in skeletal muscle following acute injury including PMNs (Ly6G^+^ cells), inflammatory monocytes (F4/80^lo^Ly6C^hi^ cells), and reparative MΦ (F4/80^hi^Ly6C^low^ cells)^76^. Ablation of ANXA1 specifically in the bone marrow compartment of healthy mice impairs myofiber regeneration following acute injury, showing that myeloid cell derived ANXA1 is indispensable for efficient skeletal muscle repair^76^. We found that endogenous ANXA1 expression was highly abundant in skeletal muscle of *mdx* mice. Interestingly, ANXA1 primarily co-localized with infiltrating immune cell populations rather than with dystrophic myofibers. Specifically, ANXA1 was predominantly expressed by interstitial PMNs (Ly6G^+^ cells) in acutely inflamed regions, by infiltrating monocytes/MΦ (CD68^+^ cells) in degenerative regions, and by interstitial M2-like MΦ (CD206^+^ cells) in regenerative regions. These localization patterns suggest that myeloid cell-derived ANXA1 may participate in multiple stages of inflammatory and regenerative responses in dystrophic muscle. In addition to infiltrating blood leukocytes, ANXA1 is expressed by resident muscle cells including endothelial cells, fibro-adipogenic progenitor cells (FAPs), and muscle stem cells (satellite cells) following acute injury^76^. Indeed, ANXA1 is known to play an important direct local role in plasma membrane repair^47^, myoblast migration^48^, and myogenic progenitor cell differentiation^49,77^. Nevertheless, the limited ANXA1 staining observed within dystrophic myofibers in the current study indicates that infiltrating blood leukocytes, rather than resident muscle cells, are likely to be a major local source of ANXA1 in DMD. This findings are consistent with an earlier study that reported that ANXA1 was highly expressed in skeletal muscle biopsy samples obtained from human DMD patients and co-localized predominantly with infiltrating CD68^+^ monocytes/MΦ^78^. ANXA1 was originally discovered as a glucocorticoid-induced secondary messenger protein that is produced by blood leukocytes^39,40,51,53^. Consistently, glucocorticoid treatment has been demonstrated to increase skeletal muscle expression mRNA expression of *Anxa1* in *mdx* mice^70^. Overall, this provides a potential cellular basis for therapies targeting the ANXA1-FPR2 pathway for the treatment of DMD, although the precise physiological role of endogenous ANXA1 in DMD remains unclear.

We found that daily systemic treatment with the ANXA1 mimetic peptide Ac2-26 in *mdx* mice markedly reduced PMN (Ly6G^+^ cell) infiltration of both the predominantly fast-twitch TA and the slow-twitch SOL, with a similar trend also observed in the left ventricle of the heart. This anti-inflammatory effect of Ac2-26 treatment was like that which was observed in mice treated daily with deflazacort in the TA, SOL, and heart. Infiltrating PMNs are known to directly contribute to secondary muscle injury in *mdx* mice^19^. Therefore, reduced intramuscular PMNs may be an important mechanism by which glucocorticoids can protect dystrophic muscle that is effectively mimicked by administration of Ac2-26. Overall, this data agrees with known actions of ANXA1–FPR2 signaling to restrict acute inflammation by both inhibiting PMN–endothelial interactions, promoting PMN apoptosis, and stimulating MΦ-mediated clearance of apoptotic PMNs^42,43^. A persistent accumulation of pro-inflammatory monocytes/MΦs can further amplify tissue damage and impair myogenic-cell proliferation in DMD^21^. In contrast, the reparative M2-like MΦ populations present in *mdx* muscle support debris clearance, myogenic differentiation, and tissue remodeling^22^. In the current study, neither deflazacort nor Ac2-26 treatment increased the absolute number of reparative M2-like MΦ (CD206^+^ cells). Nevertheless, both treatments reduced the total number of monocytes/MΦs (CD68^+^ cells) in the predominantly fast-twitch TA and thus increased the M2-to-M1 MΦ ratio in this muscle type (although this was not observed in either the SOL or heart). This suggests that Ac2-26 reproduced the effect of deflazacort to block TA muscle infiltration of pro-inflammatory blood monocytes/MΦ and/or promote their local phenotypic shift to a M2-like MΦ phenotype. This data agrees with known actions of ANXA1–FPR2 signaling to promote MΦ polarization to an anti-inflammatory/pro-regenerative subset^42,43^. However, the anti-inflammatory response to both glucocorticoids and ANXA1 mimetic peptides may depend on muscle type-specific differences in injury burden and/or fiber-type composition since neither deflazacort nor Ac2-26 was able to reduce total monocyte/MΦ infiltration of either the SOL or heart. Interestingly, Ac2-26 did uniquely appear to reduce the absolute number of M2-like MΦ (CD206^+^ cells) in the predominantly slow-twitch SOL muscle. Nevertheless, this was associated will overall beneficial rather than detrimental effects of Ac2-26 on SOL muscle histology.

Despite their overall apparently similar anti-inflammatory effects in DMD, deflazacort and Ac2-26 showed marked differences in their impact body composition and skeletal muscle growth in young *mdx* mice. Daily treatment of *mdx* mice with Ac2-26 promoted significant total body-weight gain, absolute lean mass accretion, and increased terminal hindlimb muscle weights, while simultaneously reducing the percentage of total body fat. In contrast, daily deflazacort treatment markedly reduced total body-weight gain, tended to decrease lean mass accretion, and did not positively or negatively impact terminal absolute muscle weights. Growth suppression and adverse changes in body composition are well-recognized potential consequences of long-term continuous glucocorticoid therapy^30,32,33^. The increase in absolute muscle mass in mice receiving daily Ac2-26 treatment was not equally maintained after normalization to body weight for every muscle type, indicating that the increase in muscle mass is part of a broader lean mass accretion. However, the greater absolute mass of GAST, SOL, and TA, together with increased mean TA and SOL myofiber CSA, demonstrates that skeletal muscle hypertrophy likely contributed substantially to the Ac2-26-associated increase in lean body mass. These results suggest that selectively targeting the downstream glucocorticoid-regulated ANXA1-FPR2 pro-resolving pathway may preserve beneficial control of inflammation while avoiding the well-known systemic growth-suppressive effects of direct glucocorticoid receptor activation that may result from daily steroid treatment regimens. Furthermore, lean body mass accumulation in *mdx* mice treated daily with Ac2-26 may lead to favorable metabolic adaptations including a reduction in percentage of body fat mass.

We further observed overall divergent effects of deflazacort and Ac2-26 on the contractile function of dystrophic skeletal muscle. Specifically, *mdx* mice that received daily Ac2-26 treatment generated greater *in vivo* plantar flexion torque than deflazacort-treated mice as well as greater absolute *in situ* TA force than both vehicle- and deflazacort-treated mice. However, normalization of TA contractile force to physiological muscle CSA largely eliminated these differences, and the resulting specific twitch and tetanic forces were generally similar between experimental groups. Therefore, Ac2-26 treatment appears to improve absolute muscle strength primarily by increasing skeletal muscle mass, rather than by impacting the intrinsic force-generating quality of dystrophic skeletal muscle. Our once-weekly experiments produced a similar but less pronounced pattern. Weekly Ac2-26 generally maintained greater final body weight, muscle mass, and absolute TA force than weekly deflazacort. However, once-weekly Ac2-26 treatment had no effect on lean body mass and its impact on individual muscle weights were generally more modest and inconsistent than those observed with daily treatment. These findings suggest that continuous treatment may be important for the anabolic or regenerative actions of Ac2-26. Consistent with the prior observations that intermittent glucocorticoid regimens may reduce some adverse side effects^32^, we did observe that weekly deflazacort had less suppressive effect on body-weight gain than common daily treatment regiments. Nevertheless, once-weekly deflazacort treatment of young *mdx* mice clearly still had apparently deleterious effects on absolute *in situ* TA strength in the current study when compared to mice receiving once-weekly treatment with either vehicle control or Ac2-26.

Histological analysis further supported an overall pro-regenerative effect of daily Ac2-26 administration upon dystrophic skeletal muscle. Daily treatment with both Ac2-26 and deflazacort increased type IIA, IIX, and IIB myofiber CSA in the predominantly fast-twitch TA muscle. However, only Ac2-26 increased the overall proportion of regenerating myofibers and promoted myonuclear accretion in the fast-twitch TA. Moreover, daily treatment with Ac2-26, but not deflazacort, also increased type I, IIA, and IIX myofiber CSA in the predominantly slow-twitch SOL muscle. This overall shift toward larger multinucleated fibers together with increased muscle mass, larger myofiber CSA, and greater absolute strength suggests enhanced growth and myonuclear accretion of regenerating myofibers in mice receiving daily Ac2-26 treatment. This interpretation is also supported by the direct pro-myogenic effects of Ac2-26 observed *in vitro*. In contrast, although daily deflazacort treatment did improve TA myofiber size, it did not improve lean mass or absolute force and, in fact, shifted the SOL CSA frequency distribution towards an increased proportion of smaller myofibers. These findings demonstrate that an increase in TA myofiber CSA in response to deflazacort does not necessarily reflect a generalized improvement in muscle growth or function. Despite this, both deflazacort and Ac2-26 treatment did result in a switch to a relatively faster SOL fiber-type composition (IIA → IIX). The functional significance of this fiber type shift requires further investigation given that we did not measure the contractile function of isolated SOL muscles from these *mdx* mice.

Our *in vitro* cell-culture experiments further demonstrate that Ac2-26 has direct pro-myogenic effects on dystrophic myoblasts in addition to its potent immunomodulatory actions in DMD. Previous studies had already shown that endogenous ANXA1 expression is essential for successful migration, differentiation, and fusion of healthy skeletal myoblasts in non-DMD models^48,49,77^. Consistent with these findings, we found that exogenous Ac2-26 treatment promoted myogenic differentiation and myotube formation in *D2-mdx* primary cell cultures. Interestingly, deflazacort treatment generally did not reproduce these obvious pro-myogenic effects and rather reduced the proportion of myoblasts undergoing myogenic differentiation. Therefore, the broader *in vivo* pro-regenerative benefits of Ac2-26 may result, in part, from direct stimulation of dystrophic myogenic cells in addition to facilitating a pro-regenerative immune environment. Interestingly, pharmacological inhibition of endogenous FPR2 signaling with the selective antagonist WRW4 also markedly reduced basal *in vitro* myotube formation of D2-mdx myoblasts. This pattern suggests that while infiltrating blood leukocytes appear to be the major cellular source of endogenous ANXA1 in dystrophic skeletal muscle, transcellular ANXA1-FPR2 signaling between infiltrating immune cells and resident muscle stem cells may be particularly important for myogenic-cell proliferation and fusion in DMD.

Taken together, our findings support a model in which Ac2-26 administration improves dystrophic muscle through complementary actions on both infiltrating blood leukocytes and resident myogenic cell populations. Within the myeloid compartment, Ac2-26 limits excessive PMN and inflammatory monocyte/MΦ accumulation and thereby promote a favorable local tissue environment favorable for inflammation-resolution. Within the myogenic compartment, Ac2-26 supports satellite cell accumulation, migration, and fusion. Such coordinated immunomodulatory and pro-regenerative effects would be especially advantageous in DMD, where chronic inflammation and repeated degeneration occur alongside an increasingly ineffective regenerative response. Nevertheless, other cellular targets may also contribute. For example, FPR2 is known to be expressed by multiple cell types, and the current study did not evaluate endothelial cells, fibro-adipogenic progenitors, satellite cells *in situ*, or the potential effects of Ac2-26 on systemic metabolism. Therefore, the relative contribution of immune-cell-dependent and myogenic-cell-autonomous mechanisms therefore remains to be established.

In conclusion, long-term treatment with the ANXA1 mimetic peptide Ac2-26 reduced skeletal muscle inflammation in the preclinical *mdx* mouse model of DMD to an extent comparable with deflazacort, while producing greater benefits for lean mass, muscle growth, myofiber regeneration, and absolute force production. The localization of ANXA1 to infiltrating myeloid cells and the direct stimulation of dystrophic myogenesis by Ac2-26 suggest that the ANXA1–FPR2 axis links inflammation-resolution with skeletal muscle regeneration. Although additional studies are required to establish receptor dependency *in vivo* and determine long-term safety and efficacy of Ac2-26 administration, these findings provide a rationale for further development of ANXA1 mimetic peptides as potential steroid-sparing therapies for DMD.

## ACKNOWLEDGEMENTS

The authors thank Shihuan Kuang (Duke University, formerly of Purdue University) for laboratory support and helpful discussions. The BA-D5 (developed by Schiaffino, S.), SC-71c (developed by Schiaffino, S.), BF-F3 (developed by Schiaffino, S.), MF20 (developed by Fischman, D.A.) and F5D (developed by Wright, W.E.) monoclonal antibodies were obtained from the Developmental Studies Hybridoma Bank (DSHB), created by the NICHD of the NIH and maintained at The University of Iowa, Department of Biology, Iowa City, IA 52242.

## AUTHOR CONTRIBUTIONS

**Hamood Rehman:** Conceptualization, Methodology, Software, Investigation, Formal analysis, Validation, Visualization, Writing – original draft, Writing – review & editing. **Xinyue Lu:** Investigation, Formal analysis, Writing – review & editing. **Savannah M Nasrallah:** Investigation, Formal analysis, Writing – review & editing. **Binayok Sharma:** Investigation, Formal analysis, Writing – review & editing. **Sophie Lies:** Investigation, Formal analysis, Writing – review & editing. **Shikha Adhikari:** Investigation, Formal analysis, Writing – review & editing. **Steven S. Welc:** Methodology, Formal analysis, Writing – review & editing, Supervision, Funding acquisition. **James F. Markworth:** Conceptualization, Methodology, Formal analysis, Resources, Writing – review & editing, Visualization, Supervision, Project administration, Funding acquisition. All authors approved the final draft and took responsibility for the data’s accuracy.

---

**Figure S1:**
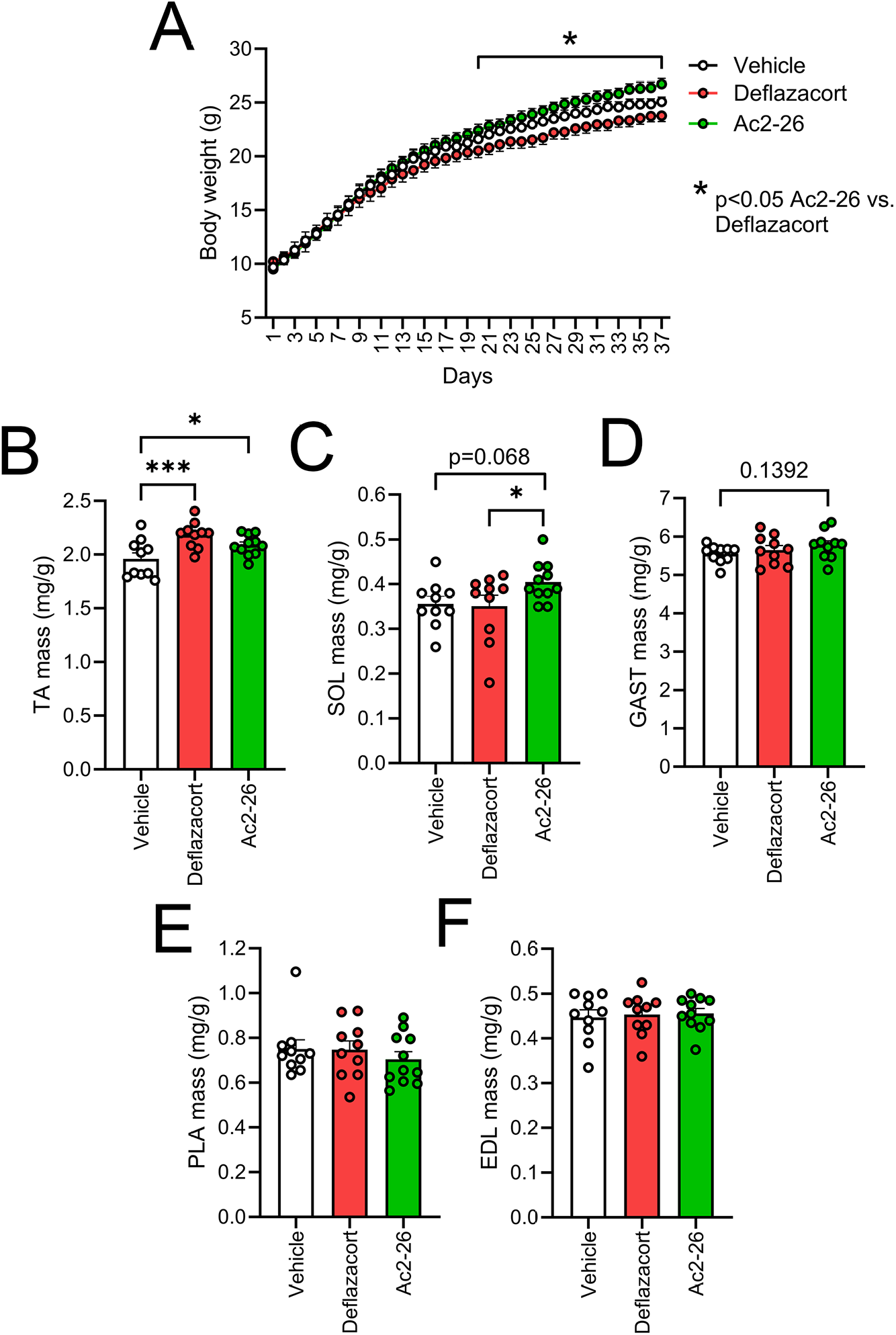
Effect of daily systemic treatment with deflazacort or Ac2-26 on absolute body weight and relative muscle mass in *mdx* mice. **A:** Longitudinal tracking of absolute body weight in male dystrophin deficient *mdx* mice that received once-daily intraperitoneal (IP) injections with vehicle control (0.25% DMSO), deflazacort (1 mg/kg/day), or Ac2-26 (1 mg/kg/day) from 3-weeks to 8-weeks of age. **B–F:** Relative weights (mg/g body weight) of isolated hindlimb muscles including tibialis anterior (TA) **(B)**, soleus (SOL) **(C)**, gastrocnemius (GAST) **(D)**, plantaris (PLA) **(E)**, and extensor digitorum longus (EDL) **(F)** obtained from 8-week-old male dystrophin deficient *mdx* mice that received once-daily intraperitoneal (IP) injections with vehicle control (0.25% DMSO), deflazacort (1 mg/kg/day), or Ac2-26 (1 mg/kg/day) for a duration of 5-weeks. Data are presented as mean ± SEM (n = 10–11 mice per group). Statistical analysis was performed by one-way ANOVA followed by Fisher’s LSD post-hoc tests. *p < 0.001, ***p < 0.001 between the indicated experimental groups.

**Figure S2:**
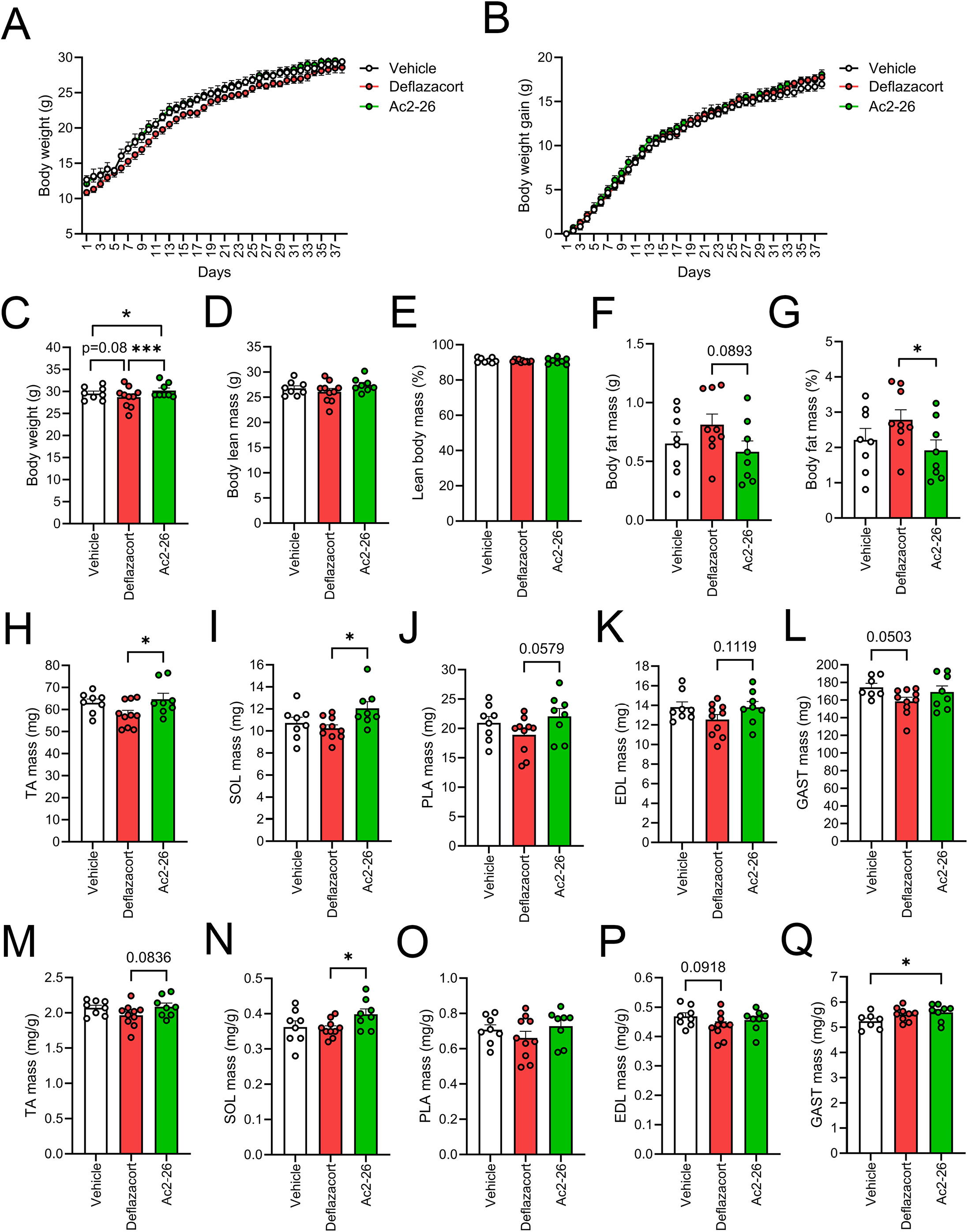
Effect of weekly systemic treatment with deflazacort or Ac2-26 on body weight, body composition, and muscle mass in *mdx* mice. **A-B:** Longitudinal tracking of absolute body weight **(A)** and cumulative body weight gain **(B)** of 8-week-old male dystrophin-deficient *mdx* mice that received once-weekly intraperitoneal (IP) injections with vehicle control (0.25% DMSO), deflazacort (1 mg/kg/week), or Ac2-26 (1 mg/kg/week) for a duration of 5-weeks. **C-G:** Endpoint EchoMRI analysis of body composition displaying final body weight **(C)**, absolute body lean mass **(D)**, percentage lean body mass **(E)**, absolute body fat mass **(F)**, and percentage body fat mass **(G)**. **H-L:** Absolute weights of isolated hindlimb muscles including tibialis anterior (TA) **(H)**, soleus (SOL) **(I)**, plantaris (PLA) **(J)**, extensor digitorum longus (EDL) **(K)**, and plantaris (PLA) **(J)**, gastrocnemius (GAST) **(L)**. **M-Q:** Relative weights (mg/g body weight) of isolated muscles including TA **(M)**, SOL **(N)**, PLA **(O)**, EDL **(P)**, GAST (**Q**). Data are presented as the mean ± SEM (n = 8–10 mice per group) with dot plots representing data from each individual mouse (biological replicates). Statistical analysis was performed by one-way ANOVA followed by Fisher’s LSD post-hoc tests. *p < 0.001, ***p < 0.001 between the indicated experimental groups.

**Figure S3:**
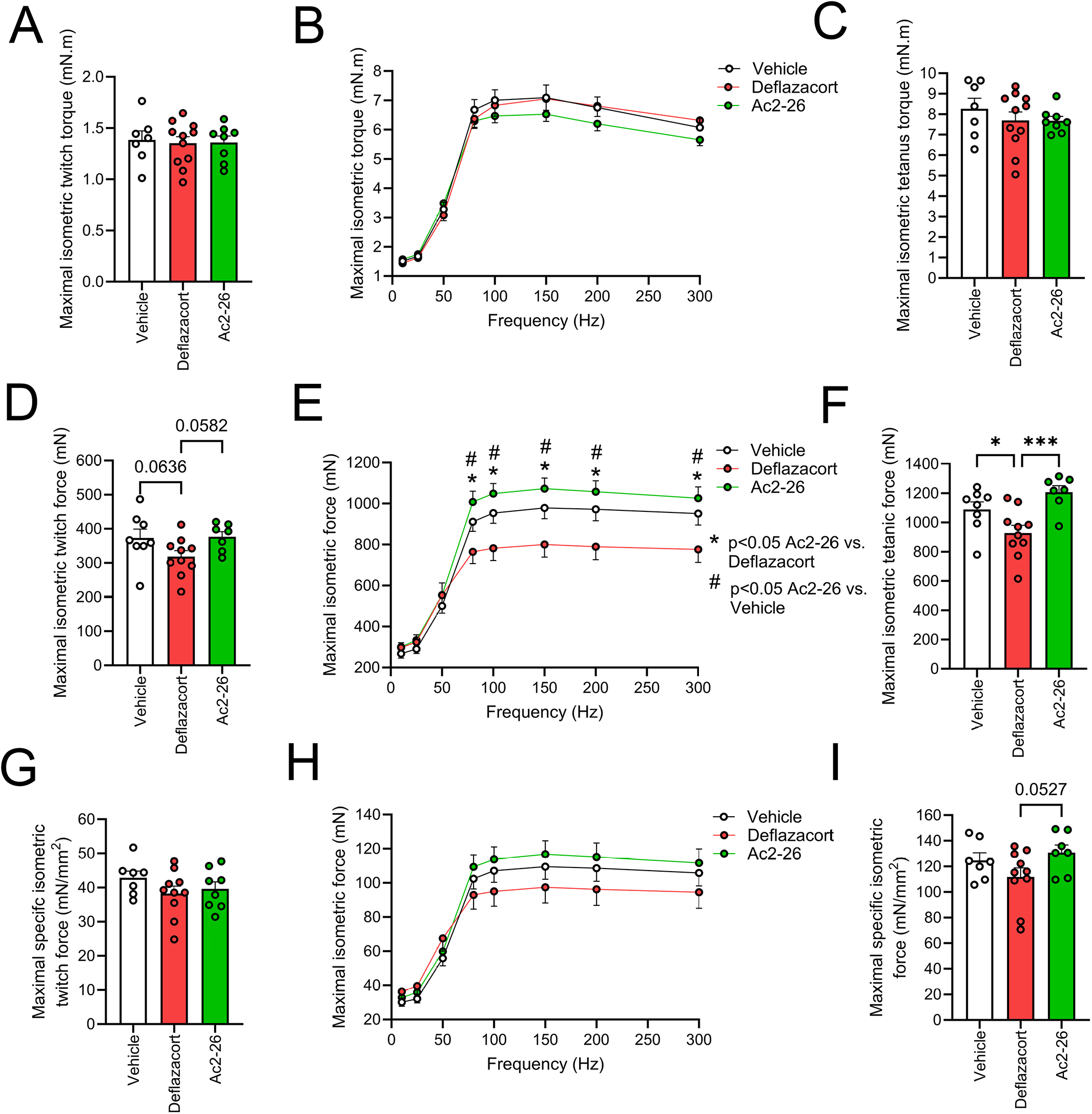
Effect of weekly systemic treatment with deflazacort or Ac2-26 on contractile function of dystrophic skeletal muscle: **A–C:** *In vivo* ankle plantar flexion isometric torque (mN-m) parameters including maximal twitch force **(A)**, the force-frequency relationship **(B)**, and maximal tetanic force **(C)** from 8-week-old male dystrophin deficient *mdx* mice that received once-weekly intraperitoneal (IP) injections with vehicle control (0.25% DMSO), deflazacort (1 mg/kg/week), or Ac2-26 (1 mg/kg/week) for a duration of 5-weeks. **D–F**: *In situ* tibialis anterior (TA) absolute isometric force (mN) parameters displaying maximal twitch force **(D)**, the force-frequency relationship **(E)**, and maximal tetanic force **(F)**. **G–I:** *In situ* normalized TA specific force (mN/mm^2^) parameters displaying maximal specific twitch force **(G)**, specific force-frequency relationship **(H)**, and maximal specific tetanic force **(I)**. **A, C, D, F, G, I:** Data are presented as the mean ± SEM (n = 8–10 mice per group) with dot plots representing data from each individual mouse (biological replicates). Statistical analysis was performed by one-way ANOVA followed by Fisher’s LSD post-hoc tests. \**p* < 0.05, \*\**p* < 0.01 between the indicated experimental groups. **B, E, H:** Data are presented as the mean ± SEM (n = 8–10 mice per group). Statistical analysis was performed by two-way mixed model ANOVA followed by Fisher’s LSD post-hoc tests. \**p* < 0.05 Ac2-26 vs. deflazacort group; #*p* < 0.05 Ac2-26 vs. vehicle group.

**Figure S4:**
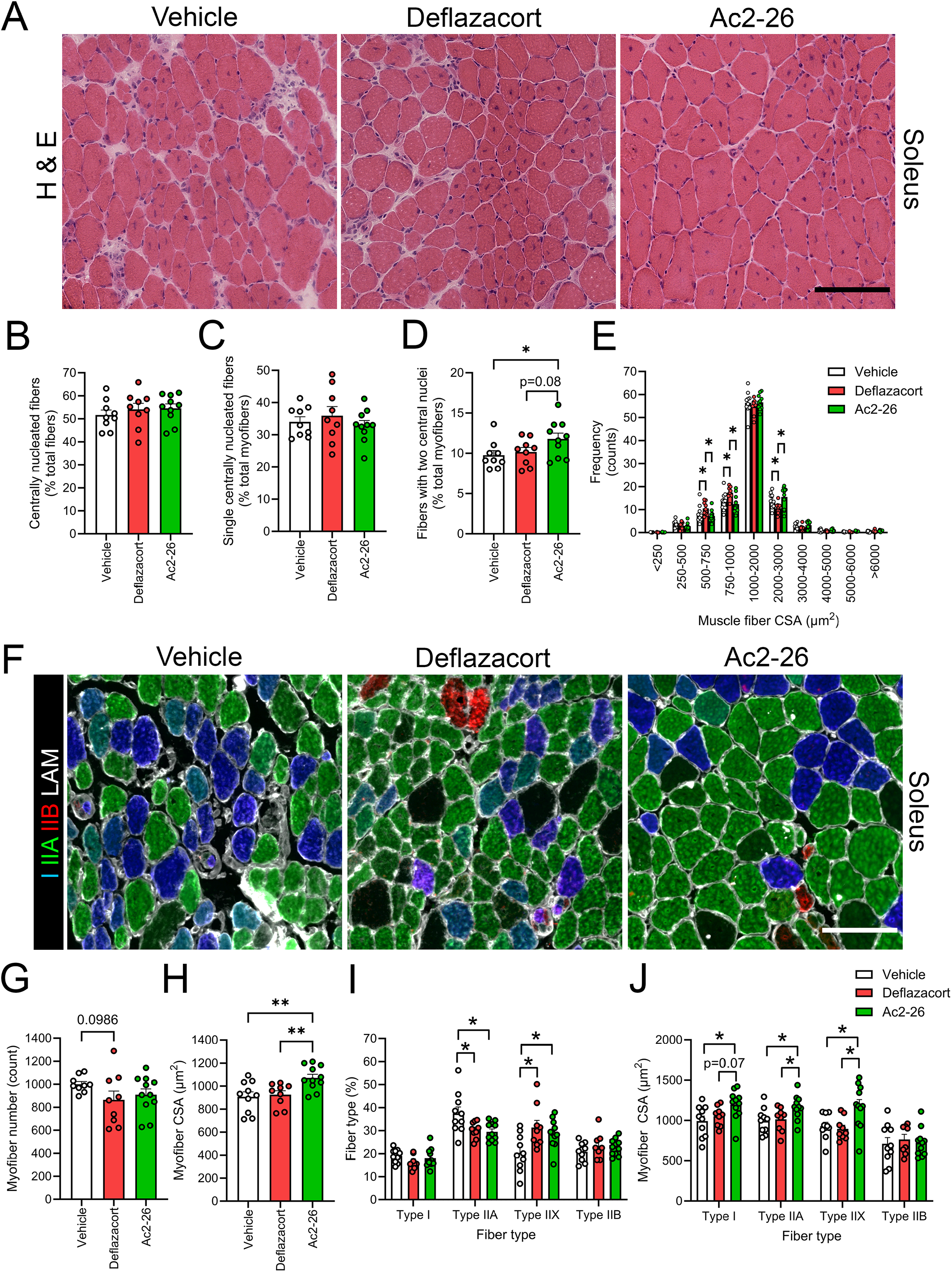
Histological and immunofluorescence analysis of slow-twitch soleus myofiber regeneration and muscle fiber type composition of *mdx* mice receiving daily systemic treatment with deflazacort or Ac2-26. **A:** Representative hematoxylin and eosin (H & E) stained soleus (SOL) muscle cross-sections from 8-week-old dystrophin deficient male *mdx* mice that received once-daily intraperitoneal (IP) injections with vehicle control (0.25% DMSO), deflazacort (1 mg/kg/day), or Ac2-26 (1 mg/kg/day) for a duration of 5-weeks. Quantification of the total proportion of regenerating (centrally nucleated) myofibers **(B)**, percentage of single centrally nucleated fibers **(C)**, percentage of fibers with two central nuclei **(D)**, and frequency distribution of muscle fiber cross-sectional area (CSA) **(E)**. **F:** Representative immunofluorescence images of SOL muscle cross-sections stained with primary antibodies against for type I myosin heavy chain (MyHC; blue), type IIA MyHC (green), type IIB (red), and laminin (LAM; white). Type IIX fibers (black) are identified by lack of positive staining for type I, IIA, and IIB MyHC. **G–J:** Quantitative analysis of total SOL myofiber number **(G)**, overall mean myofiber CSA **(H)**, fiber type percentage **(I)**, and fiber type-specific myofiber CSA **(J)**. Data are presented as mean ± SEM (n = 10–11 mice per group) with dot plots representing data from each individual mouse (biological replicates). Statistical analysis was performed using a one-way ANOVA followed by Fisher’s LSD post-hoc tests. \**p* < 0.05, \*\**p* < 0.01, \*\*\**p* < 0.001 between the indicated experimental groups. Scale bars are 100 µm.

**Figure S5:**
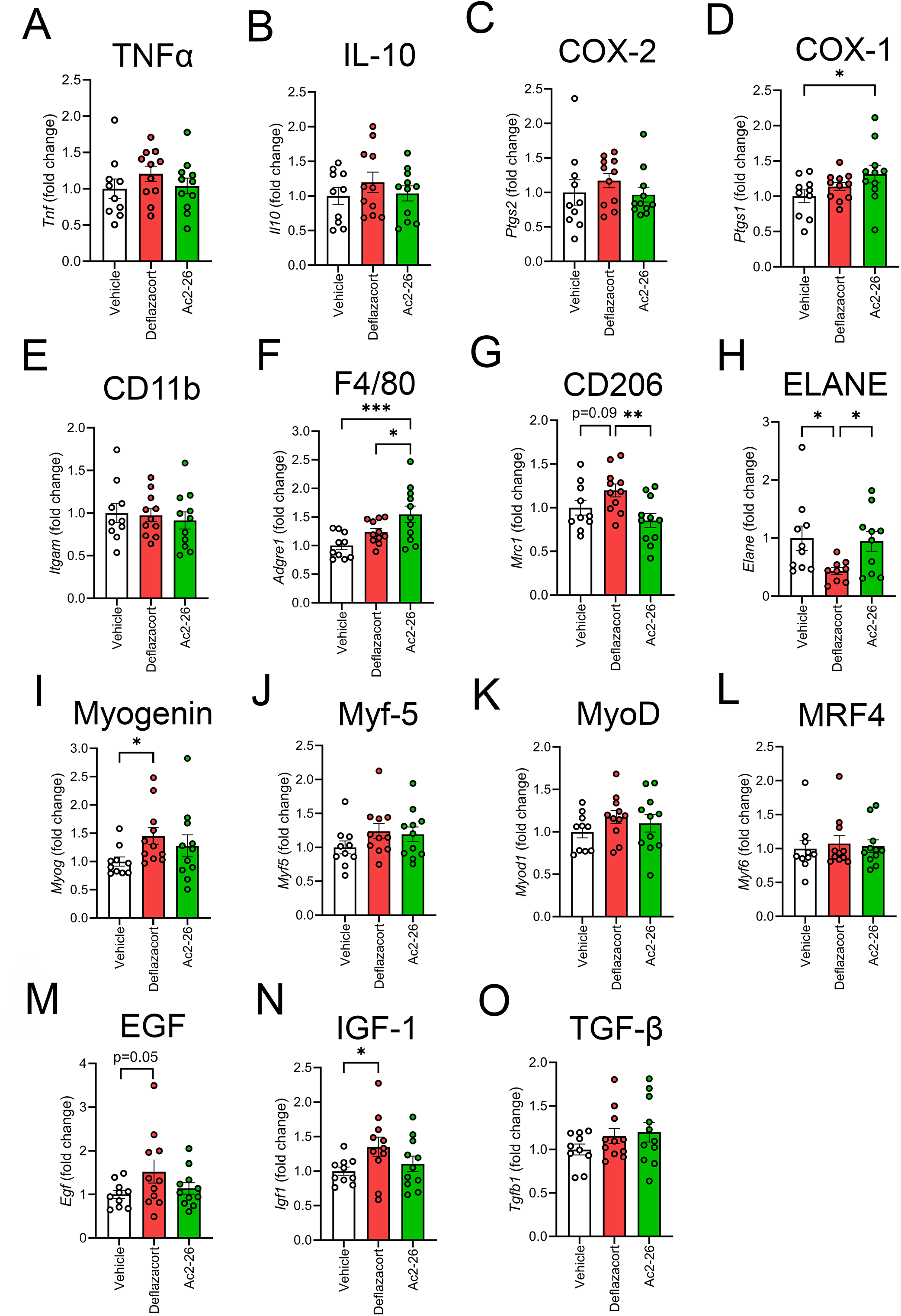
Dystrophic skeletal muscle gene expression following daily systemic administration of deflazacort or Ac2-26. Gastrocnemius (GAST) muscles were obtained from 8-week-old male *mdx* mice that had received once-daily intraperitoneal (IP) injections with vehicle control (0.25% DMSO), deflazacort (1 mg/kg/day), or Ac2-26 (1 mg/kg/day) for a duration of 5-weeks. Total RNA was extracted and gene expression analyzed by reverse transcription-quantitative polymerase chain reaction (RT-qPCR). **A–D:** GAST mRNA expression of inflammation-related genes including tumor necrosis factor alpha (TNF-α, *Tnf*) **(A)**, interleukin-10 (IL-10, *Il10*) **(B)**, cyclooxygenase-2 (COX-2, *Ptgs2*) **(C)**; and cyclooxygenase-1 (COX-1, *Ptgs1*) **(D)**. **E–H:** GAST mRNA expression of immune cell markers including integrin alpha M (CD11b, *Itgam*) **(E)**, adhesion G protein-coupled receptor E1 (F4/80, *Adgre1*) **(F)**, mannose receptor C-type 1 (CD206, *Mrc1*) **(G)**, and neutrophil elastase (NE, *Elane*) **(H)**. **I–L:** GAST mRNA expression of myogenic regulatory factors including myogenin (MyoG, *Myog*) (**I**), myogenic factor 5 (Myf-5, *Myf5*) **(J)**, myogenic differentiation protein 1 (MyoD1, *Myod1*) (**K**), myogenic regulatory factor 4 (MRF4, *Myf6*) **(L)**. **M–O:** GAST mRNA expression of growth factors including epidermal growth factor (EGF, *Egf*) **(M)**, insulin-like growth factor 1 (IGF-1, *Igf1*) **(N)**, and transforming growth factor beta 1 (TGF-β1, *Tgfb1*) **(O)**. Data are presented as the group mean ± SEM (n = 10–11 mice per group). Statistical analysis was performed by one-way ANOVA followed by Fisher’s LSD post hoc test. \**p* < 0.05, \*\**p* < 0.01, and \*\*\**p* < 0.001 between the indicated experimental groups.

**Figure S6:**
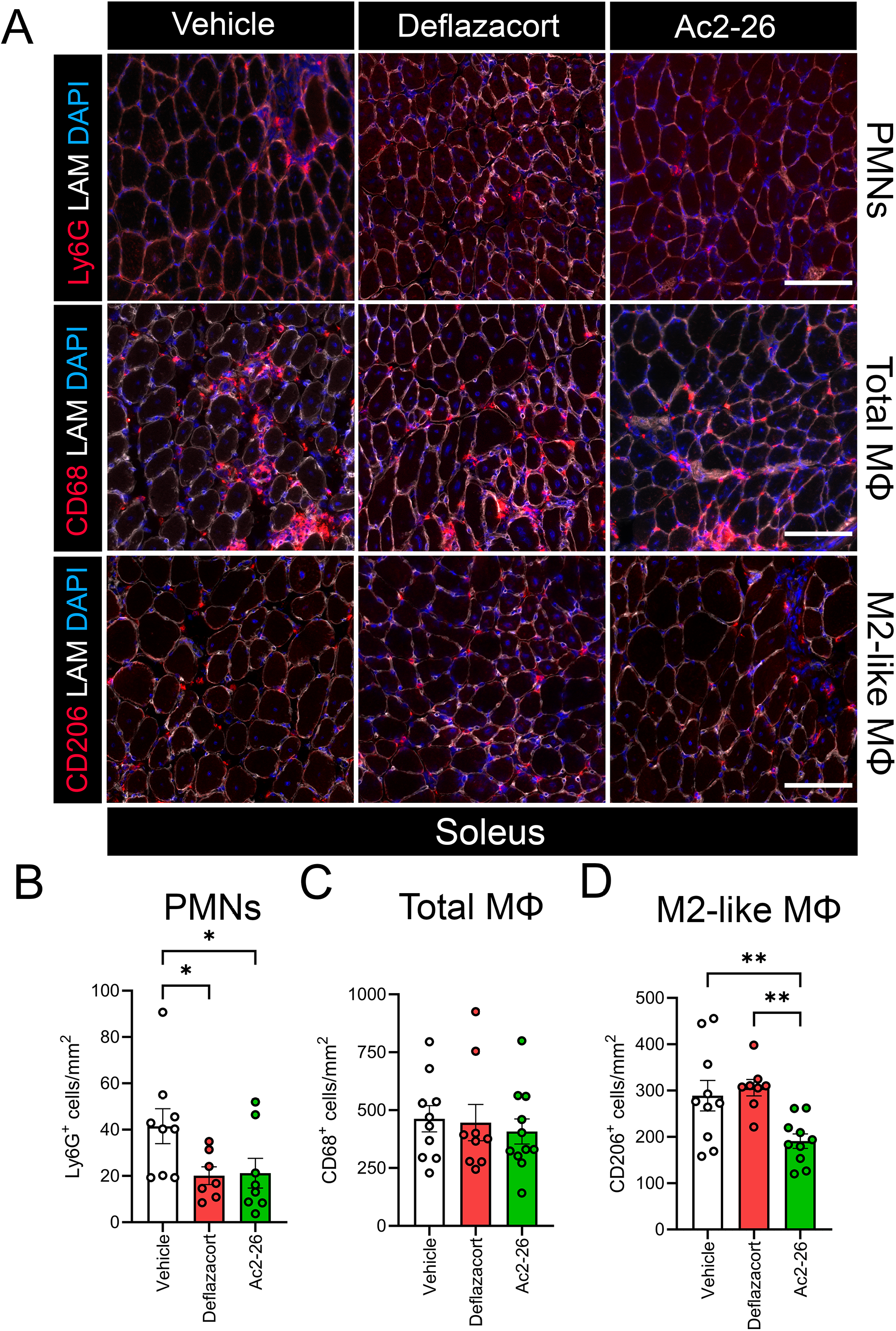
Immunofluorescence analysis of infiltrating immune cell populations in slow-twitch soleus muscles of *mdx* mice following daily systemic deflazacort or Ac2-26 administration. **A:** Representative immunofluorescent images of soleus (SOL) muscle cross-sections from 8-week-old male dystrophin deficient *mdx* mice that received once-daily intraperitoneal (IP) injections with vehicle control (0.25% DMSO), deflazacort (1 mg/kg/day), or Ac2-26 (1 mg/kg/day) for a duration of 5-weeks were stained for distinct immune cell markers (red), laminin (LAM; white), and cell nuclei (DAPI; blue). Rows from top to bottom show neutrophils (Ly6G^+^ cells), total monocytes/macrophages (CD68^+^ cells), and M2-like macrophages (CD206^+^ cells) across the treatment groups**. B–D:** Quantitative analysis of intramuscular numbers of neutrophils (Ly6G^+^ cells) **(B)**, total monocytes/macrophages (CD68^+^ cells) **(C)**, and the M2/M1 macrophage ratio **(D).** Data are presented as mean ± SEM (n = 10–11 mice per group) with dot plots representing data from each individual mouse (biological replicates). Statistical analysis was performed by one-way ANOVA followed by Fisher’s LSD post-hoc tests. \**p* < 0.05, \*\**p* < 0.01, \*\*\**p* < 0.001 between the indicated experimental groups. Scale bars are 100 µm.

**Figure S7:**
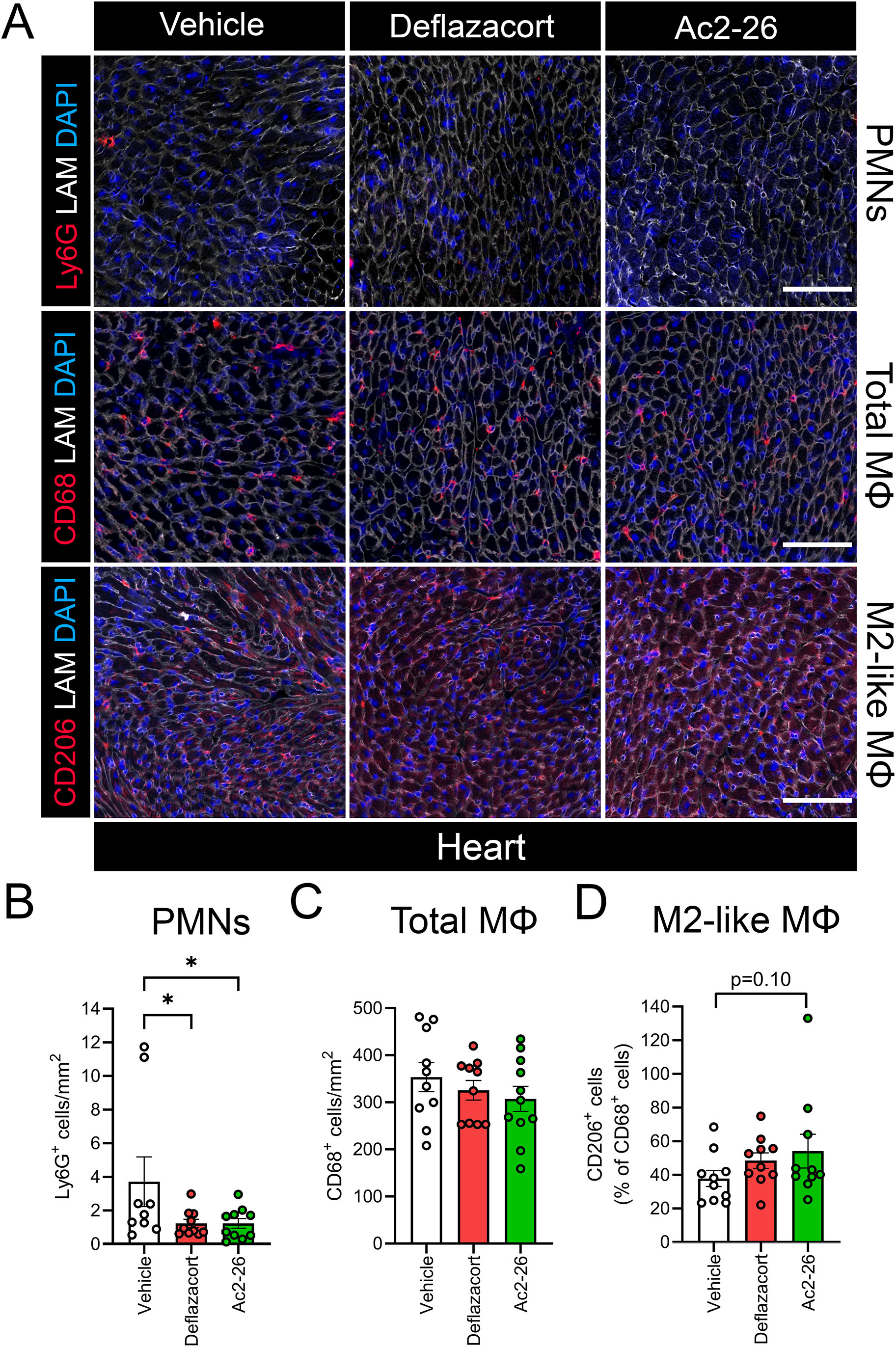
Immunofluorescence analysis of infiltrating immune cell populations in the heart of *mdx* mice following daily systemic deflazacort or Ac2-26 administration. **A:** Representative immunofluorescent images of cross-sections obtained from the left ventricle of the heart of 8-week-old male dystrophin deficient *mdx* mice that received once-daily intraperitoneal (IP) injections with vehicle control (0.25% DMSO), deflazacort (1 mg/kg/day), or Ac2-26 (1 mg/kg/day) for a duration of 5-weeks were stained for distinct immune cell markers (red), laminin (LAM; white), and cell nuclei (DAPI; blue). Rows from top to bottom show neutrophils (Ly6G^+^ cells; red), total monocytes/macrophages (CD68^+^ cells; red), and M2-like macrophages (CD206^+^ cells; red) across the treatment groups**. B–D:** Quantitative analysis of intramuscular numbers of neutrophils (Ly6G^+^ cells) **(B)**, total monocytes/macrophages (CD68^+^ cells) **(C)**, and the M2/M1 macrophage ratio **(D).** Data are presented as mean ± SEM (n = 10–11 mice per group) with dot plots representing data from each individual mouse (biological replicates). Statistical analysis was performed by one-way ANOVA followed by Fisher’s LSD post-hoc tests. Scale bars are 100 µm. \**p* < 0.05 between the indicated experimental groups.

